# Unravelling a novel role for Cannabidivarin in the modulation of subventricular zone postnatal neurogenesis

**DOI:** 10.1101/2023.05.16.540990

**Authors:** Diogo M. Lourenço, Rita Soares, Sónia Sá Santos, Joana M. Mateus, Rui S. Rodrigues, João B. Moreira, Sandra H. Vaz, Ana M. Sebastião, Susana Solá, Sara Xapelli

**Author notes:** Correspondence Sara Xapelli, Instituto de Farmacologia e Neurociências, Faculdade de Medicina, Universidade de Lisboa and Instituto de Medicina Molecular João Lobo Antunes, Faculdade de Medicina, Universidade de Lisboa, 1649-028 Lisboa, Portugal. **Conflict of Interest:** The authors declare no competing interests.

## Abstract

Postnatal neurogenesis has been shown to rely on the endocannabinoid system. Here we aimed at unravelling the role of Cannabidivarin (CBDV), a non-psychoactive cannabinoid, with high affinity for the non-classical cannabinoid receptor TRPV1, on subventricular zone (SVZ) postnatal neurogenesis. Using the neurosphere assay, SVZ-derived neural stem/progenitor cells (NSPCs) were incubated with CBDV and/or 5’-Iodoresinferotoxin (TRPV1 antagonist), and their role on cell viability, proliferation, and differentiation were dissected. CBDV was able to promote, through a TRPV1-dependent mechanism, cell survival, cell proliferation and neuronal differentiation. Furthermore, pulse-chase experiments revealed that CBDV-induced neuronal differentiation was a result of cell cycle exit of NSPCs. Regarding oligodendrocyte differentiation, CBDV inhibited oligodendrocyte differentiation and maturation. Since our data suggested that the CBDV-induced modulation of NSPCs acted via TRPV1, a sodium-calcium channel, and that intracellular calcium levels are known regulators of NSPCs fate and neuronal maturation, single cell calcium imaging was performed to evaluate the functional response of SVZ-derived cells. We observed that CBDV-responsive cells displayed a two-phase calcium influx profile, being the initial phase dependent on TRPV1 activation. Taken together, this work unveiled a novel and untapped neurogenic potential of CBDV via TRPV1 modulation. These findings pave the way to future neural stem cell biological studies and repair strategies by repurposing this non-psychoactive cannabinoid as a valuable therapeutic target.

## 1. Introduction

Adult neural stem/progenitor cells (NSPCs) can be found in the subventricular zone, (SVZ) lining the lateral walls of the lateral ventricles (Bond *et al*. 2015). Importantly, adult NSPCs have self-renewing properties and are tri-potent, having the capacity to generate neurons, astrocytes and oligodendrocytes (Soares *et al*. 2020). In the SVZ, besides originating neuroblasts (Luskin and Boone 1994), a minority of NSPCs enter the oligodendroglial lineage (Suzuki and Goldman 2003; Menn *et al*. 2006) by originating oligodendrocyte progenitor cells (OPCs) that migrate into the surrounding cortex and white matter, differentiating into myelinating oligodendrocytes (Aguirre and Gallo 2004; Butti *et al*. 2019).

Adult NSPCs are mostly quiescent *in vivo* (Cavallucci *et al*. 2016). The balance between quiescence and activity regulates, not only the rate of cytogenesis, but also the long-term maintenance of the NSPC pool (Urbán *et al*. 2019; Cheung and Rando 2013). Therefore, finding and exploiting modulators of NSPCs that can boost postnatal neurogenesis and oligodendrogenesis is imperative.

Several studies have shown that neurogenesis is heavily modulated by the endocannabinoid system, (Rodrigues *et al*. 2017; Galve-Roperh *et al*. 2013; Zimmermann *et al*. 2018; Xapelli *et al*. 2013; Molina-Holgado *et al*. 2007; Ferreira *et al*. 2018; Bravo-Ferrer *et al*. 2017). A growing body of scientific and clinical data has been attesting the potential of medical-cannabis to ameliorate symptoms of several pathologies (Pacher *et al*. 2006; Pertwee 2005; Di Marzo and De Petrocellis 2006; Schlag *et al*. 2021; Hall and Degenhardt 2009; Castaneto *et al*. 2014; Ebbert *et al*. 2018). Notwithstanding, the chronic and abusive consumption of *Cannabis sp.* has also been associated with detrimental health effects, such as cognitive and memory impairments (Cohen *et al*. 2020; Figueiredo *et al*. 2020). Thus, one of the major challenges of cannabis research is to find ways to prevent the negative side-effects associated with cannabis-based medicines (Hall and Degenhardt 2009; Grant *et al*. 2018; Connor *et al*. 2021). One possible alternative to overcome this issue can be related with the modulation of the non-classical cannabinoid receptors, such as the transient receptor potential cation channel subfamily V member 1 (TRPV1). This sodium-calcium ion channel receptor, belongs to the endovanilloid system and is associated with thermoregulation and nociception (Tominaga and Tominaga 2005; Caterina *et al*. 1997). Importantly, several studies demonstrated that TRPV1 is a potential target for the regulation of cell proliferation and apoptosis (Stock *et al*. 2014; Kong *et al*. 2010; Czaja *et al*. 2008). Previous works have highlighted that, in both mouse and rat models, TRPV1-activation was shown to promote cell death, inhibited of cell proliferation and impaired neuronal maturation (Czaja *et al*. 2008; Kong *et al*. 2010). Accordingly, TRPV1 knockout (KO) mice present higher levels of NSPC proliferation when compared to wild type (Stock *et al*. 2014). In contrast, other works reported a TRPV1-dependent increase of neuronal differentiation and promotion of axonal and neurite growth in dorsal root ganglia cultures (Frey *et al*. 2018; Goswami *et al*. 2007). TRPV1 can also be activated by cannabidivarin (CBDV), a phytocannabinoid that has negligible affinity for cannabinoid type 1 and 2 receptors (CB1R and CB2R) (Rosenthaler *et al*. 2014; Hill *et al*. 2013). CBDV has also been described as an agonist of TRPV2 and TRPA1 (De Petrocellis *et al*. 2011), as an inverse agonist of GPR6 (Laun *et al*. 2019) as well as an allosteric antagonist of GPR55 (Anavi-Goffer *et al*. 2012). However, there is a broad consensus in the literature suggesting that its biological activity is mostly via a TRPV1-dependent mechanism of action (Iannotti *et al*. 2014; Straiker *et al*. 2021; Huizenga *et al*. 2019; Thornton *et al*. 2020; Muller *et al*. 2019).

Given the role of TRPV1 in regulating cell death, cell proliferation and neuronal differentiation and, that CBDV is a non-psychoactive cannabinoid, it is relevant to thoroughly understand how this emerging cannabinoid can modulate neurogenesis and how it could be used as a viable drug for brain repair strategies. Therefore, with this work we aimed at unravelling how CBDV modulate NSPCs fate and whether those effects are mediated by TRPV1.

## 2. Material and methods

### 2.1. Ethical statement

All experiments followed the European Community (86/609/EEC; 2010/63/EU; 2012/707/EU) and Portuguese (DL 113/2013) legislation for the protection of animals used for scientific purposes. The protocol was approved by the “iMM’s institutional Animal Welfare Body – ORBEA-iMM and the National competent authority – DGAV (Direcção Geral de Alimentação e Veterinária).”

### 2.2. Animals

C57BL/6J females were kept in standard housing, grouped in pairs, while males were single housed. All animals were kept on a 12h light/dark cycle, with food and water provided *ad libitum*. No breeding attempts were made before sexual maturity was reached, at 8 weeks of age. Breeding trios were used until one-year old of age or 10 successful mating sessions, in order to maximise breeding success.

All efforts were made to minimize animal suffering and stress, and to use the minimum number of animals, according to standard and ethical procedures. All animals were given access to hiding places, in the form of disposable igloos or cardboard tubes, as well as proper nesting material and wooden pellets for chewing.

Experiments were performed *ex-vivo* with biological material obtained from postnatal day 1 to 3 (P1-3) C57BL/6J mice and subsequently maintained in *in vitro* conditions. A minimum of 3 pups was required to perform one SVZ cell culture and a minimum of 3 independent cultures was required to perform statistical analysis. All pups per litter were used per SVZ cell culture.

### 2.3. SVZ Cell Cultures

SVZ Neurospheres (3D clusters of clones of NSPCs) were prepared from early postnatal (P1-3) C57BL/6J mice as previously described (Soares *et al*. 2020). In brief, P1-3 C57BL/6J mice were decapitated with a single incision with sharp scissors at the base of the brainstem. SVZ fragments were dissected out from 450μm-thick coronal brain slices. All collected tissue was pooled and digested with 0.05% Trypsin-EDTA (#25300054, Gibco™) in Hanks’ Balanced Salt Solution (HBSS, #14175095, Gibco™) and mechanically dissociated with a P1000 pipette. Single-cell suspension was resuspended in serum-free medium (SFM), composed of Dulbecco’s Modified Eagle Medium/Nutrient Mixture F-12 + GlutaMAX™ supplement (DMEM+GlutaMAX™, #31331028, Gibco™) supplemented with 100U/mL penicillin and 100μg/mL streptomycin (#15070063, Gibco™), 1% B-27™ (#17504044, Gibco™) and growth factors (10ng/mL epidermal growth factor (EGF, #PHG0311, Gibco™) and 5ng/mL bovine fibroblast growth factor-2 (bFGF-2, #13256-029, Gibco™)) (proliferative conditions). SVZ cells were seeded in 60mm diameter Petri dishes (#430166, Corning) and maintained for six days in a 95% air/5% CO2 humified atmosphere at 37°C. Resulting neurospheres were seeded for 24h onto 12mm ⌀ glass coverslips (#631-1577P, VWR) coated with 100μg/mL poly-D-lysine (PDL, #P7886, Sigma-Aldrich) in SFM devoid of growth factors (differentiative conditions). After 24h (day 0), the medium was renewed with or without (control) a range of pharmacological treatments for 48h or 7 days.

### 2.4. Pharmacological treatments and experimental setting

Since CBDV has very weak affinity for both CB1R and CB2R (Rosenthaler *et al*. 2014) and it has been proposed to activate TRPV1 (Iannotti *et al*. 2014), we evaluated its effects on SVZ-derived NSPC survival, proliferation and differentiation and studied TRPV1-dependency using a selective TRPV1 antagonist (5’-Iodoresiniferatoxin, 5’-IRTX).

Plated neurospheres were exposed to three increasing concentrations of CBDV (drug concentration-response studies) (Table 1). These were established based on previous studies with cannabinoids (Rodrigues *et al*. 2017; Xapelli *et al*. 2013; Stanslowsky *et al*. 2017; Compagnucci *et al*. 2013) since the Ki for CBDV for TRPV1 is not known. Additionally, to study if the effect seen by CBDV was TRPV1-dependent, the TRPV1 antagonist 5’-IRTX was used. This drug is a potent vanilloid receptor antagonist, 40-fold more potent than the prototypical TRPV1 antagonist capsazepine (Wahl *et al*. 2001). Whenever cells needed to be co-treated with the antagonist (at 300nM), they were incubated 30 minutes prior to the treatment with CBDV. All drugs were dissolved in Dimethyl sulfoxide (DMSO, #D2650, Merck Life Sciences) at a stock solution of [50mM] CBDV and [10mM] 5’-IRTX. Fresh dilutions were prepared on the day of the pharmacological tests with SFM-DMEM+GlutaMAX™ devoid of growth factors.

**Table 1.** Pharmacological treatments

| Drug | Purity | Biological Activity | Concentration used | Ki for CB1R/ CB2R | Ki for TRPV1 | Company | # | References |
| --- | --- | --- | --- | --- | --- | --- | --- | --- |
| **Cannabidivarin**<br>2-((1S,6S)-3-methyl-6-(prop-1-en-2-yl)cyclohex-2-enyl)-5-propylbenzene-1,3-diol | ≥98% | TRPV1 agonist and GPR55 antagonist | 100nM<br>300nM<br>1μM | CB1R: 14711nM<br>CB2R: 574.2nM | Not Known | THC Pharm GmbH The Health Concept | - | (Anavi-Goffer *et al.* 2012; Iannotti *et al.* 2014; Rosenthaler *et al.* 2014) |
| **5'-Iodoresiniferatoxin**<br>6,7-Deepoxy-6,7-didehydro-5-deoxy-21-dephenyl-21-(phenyl Imethyl)-daphnetoxin, 20-(4-hydroxy-5-iodo-3-methoxybenzene acetate) | >99% | TRPV1 antagonist | 300nM | Not applicable | 5.8nM | Alomone Labs | I-800 | (Wahl *et al.* 2001) |

### 2.5. Total RNA isolation and quantitative real-time reverse transcription polymerase chain reaction (qRT-PCR)

Total RNA was isolated from DIV7 SVZ-derived cells, subjected to the pharmacological treatments mentioned above, using TRIzol™ Reagent (#15596026, Invitrogen™) (Vilain *et al*. 2012). Cells were scraped into a tube containing 1mL TRIzol™ and manually dissociated with a P1000 pipette After mixing with 200µL of chloroform, and vortexing for 15 seconds, a centrifugation at 12,000 × g for 10 minutes at 4°C was performed to collect the aqueous phase, to which equal volume of isopropyl alcohol was added. After a centrifugation at 17,500 × g, for 10 minutes at 4°C, RNA pellet was washed in sequential cycles of decreasing volumes of 75% ethanol (400µL–100μL). RNA pellet was air dried, for 15 minutes at room temperature (RT) and resuspended in 10μL of nuclease-free water followed by an incubation for 10 minutes at 55°C. RNA purity and concentration were obtained using Nanodrop 2000 Spectrophotometer (NanoDrop Technologies LLC). DNA contaminations were eliminated with DNase I recombinant (#04716728001; Roche Applied Science) following the manufacturer’s instructions. All samples were stored at −80°C until use.

cDNA was prepared from 1000ng total RNA using NZY Reverse Transcriptase (#MB12402; NZYTech) according to manufacturer’s instructions. Real-time RT-PCR was performed using a SensiFastTM SYBR® Hi-ROX kit (#BIO-92020; Bioline USA Inc.) in an Applied Biosystems QuantStudio 7 Flex Real-Time PCR system (Thermo Fisher Scientific Inc.). Primer sequences are listed in Table 2. Relative gene expression was calculated based on the standard curve and normalized to the level of hypoxanthine glyceraldehyde 3-phosphate dehydrogenase (GAPDH) housekeeping gene and expressed as fold change from controls. Primers were designed using the Reference Genome GRCm39 (Ensembl Assembly Genome C57BL_6NJ_v1; Accession: GCA_001632555.1) taken from the Ensembl Release 100 (April 2020).

**Table 2.**
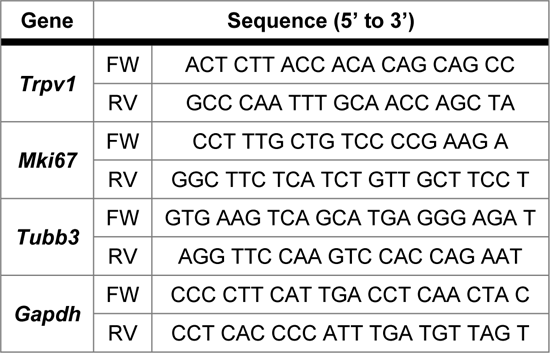
Primers

### 2.6. Evaluation of Cell viability, Cell proliferation and Cell differentiation under differentiative conditions and immunocytochemistry (ICC) assays

#### 2.6.1. Cell viability

To investigate the effect of the different pharmacological treatments on cell viability, SVZ cells were exposed, at DIV2, to 3μg/mL of propidium iodide (PI, # P4170, Sigma-Aldrich) for 30min before fixation. PI is a fluorescent intercalating agent that binds to DNA in cells that have a compromised cell membrane, thus is useful to differentiate healthy cells from necrotic or late apoptotic cells (Lecoeur 2002). Cells were processed for ICC, as mentioned in 2.5.5.

#### 2.6.2. Cell proliferation

Cell proliferation was assessed at DIV2 by co-incubating cells with 10μM of 5-bromo-2’-deoxyuridine (BrdU) (#B5002, Sigma-Aldrich) in the last 4h of the pharmacological treatments. BrdU is a synthetic thymidine analogue able to substitute thymidine in the DNA double chain during the S Phase of the cell cycle (Kee *et al*. 2002). Cells were prepared for ICC, as mentioned in 2.5.5. BrdU was unmasked by permeabilizing cells in PBS with 1% Triton™ X-100 at RT for 30min. DNA was denatured in 1M HCl for 20min at 37°C. See antibodies on Table 3.

**Table 3.**
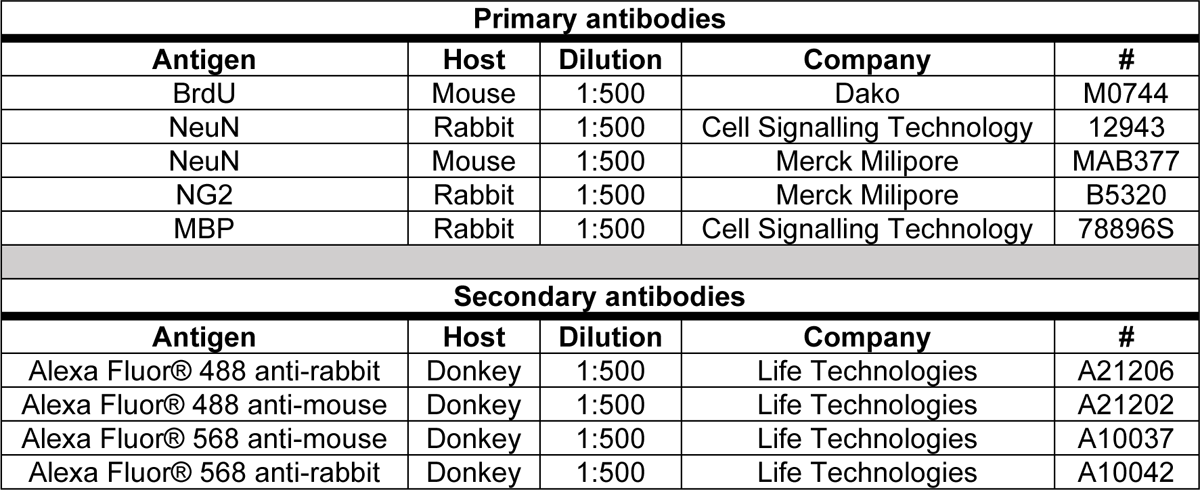
Primary and Secondary antibodies

| Primary antibodies |  |  |  |  |
| --- | --- | --- | --- | --- |
| Antigen | Host | Dilution | Company | # |
| BrdU | Mouse | 1:500 | Dako | M0744 |
| NeuN | Rabbit | 1:500 | Cell Signalling Technology | 12943 |
| NeuN | Mouse | 1:500 | Merck Milipore | MAB377 |
| NG2 | Rabbit | 1:500 | Merck Milipore | B5320 |
| MBP | Rabbit | 1:500 | Cell Signalling Technology | 78896S |
| Secondary antibodies |  |  |  |  |
| Antigen | Host | Dilution | Company | # |
| Alexa Fluor® 488 anti-rabbit | Donkey | 1:500 | Life Technologies | A21206 |
| Alexa Fluor® 488 anti-mouse | Donkey | 1:500 | Life Technologies | A21202 |
| Alexa Fluor® 568 anti-mouse | Donkey | 1:500 | Life Technologies | A10037 |
| Alexa Fluor® 568 anti-rabbit | Donkey | 1:500 | Life Technologies | A10042 |

#### 2.6.3. Cell differentiation

To assess cell differentiation, cells were fixed at two different timepoints, DIV2 and DIV7, and handled for ICC, as mentioned in 2.5.5. Neuronal and oligodendroglial lineages were evaluated using the respective antibodies on Table 3.

#### 2.6.4. Pulse-chase experiments

SVZ-derived cells were given a BrdU pulse for the first 24h of drug treatments followed by a chase of 6 days (without BrdU) in the absence (control) or presence of our pharmacological treatments. 7-day treated SVZ-derived cells, were fixed and prepared for ICC according to the protocol described in 2.5.5. For antibodies see Table 3.

#### 2.6.5. ICC assay

Cells were fixed in 4% PFA for 30 minutes and washed with PBS at RT. An incubation with PBS with 0.5% Triton X-100 (#X100, Sigma-Aldrich) and 3% bovine serum albumin (BSA, #MB04602, NZYTech) for 1.5h was performed to block nonspecific binding sites, followed by an overnight incubation with the primary antibodies (Table 3). The next day, after several washes with PBS, the corresponding secondary antibodies (Table 3) were incubated for 1h at RT, followed by nuclei counterstaining with Hoechst 33342 (12μg/mL in PBS, #H1399, Invitrogen™) and mounting in Mowiol fluorescent medium (#324590, Sigma-Aldrich).

### 2.7. Evaluation of cell proliferation under proliferative conditions

SVZ neurospheres were prepared according to the protocol mentioned in 2.2 by growing SVZ cells in proliferative conditions (SFM supplemented with EGF and bFGF-2) in the presence or absence (control condition) of CBDV (1µM). SVZ-derived cells were allowed to form neurospheres and the analysis of proliferation occurred after 5 days in culture. Neurosphere size (diameter, DN, and area, AN,) was used as an indirect indicator of cell proliferation (Mori *et al*. 2006).

### 2.8. Microscopy

Immunofluorescence images were captured using an AxioCamMR3 monochrome digital camera (Carl Zeiss Inc.) mounted on a Zeiss Axiovert 200M inverted widefield fluorescence microscope (Carl Zeiss Inc.), with a 40x EC Plan-NeoFluar (NA 0.75) objective. Images were obtained using the software AxioVision 4 (Carl Zeiss Inc.), stored and analysed in an uncompressed 8-bit Tiff (.tiff) format. One pixel corresponds to 0.25μm and the captured image size was 1388 x 1040 pixels.

Phase contrast images of neurospheres were captured using an AxioCam 208c colour digital camera (Carl Zeiss Inc.) mounted on a Zeiss Primovert inverted widefield microscope (Carl Zeiss Inc.), with a 4x Plan Achromat (NA 0.1) Ph0 objective. Images were obtained using the software Zen 3.2 (Blue edition) (Carl Zeiss Inc.), stored and analysed in an uncompressed Carl Zeiss Image (.czi) format. One pixel corresponds to 0.925μm and the captured image size was 3840 x 2160 pixels.

### 2.9. Calcium imaging

SVZ-derived cells were cultured as mentioned in 2.2 and cells were seeded in μ-ibidi 8 well plates (#80826, ibidi). The Ca^2+^ imaging assay was performed in SVZ cells 7 days after seeding using a Zeiss Axiovert 135TV inverted microscope with epifluorescent optics and equipped with a high-speed multiple excitation fluorimetric system (Lambda DG4, with a 175W Xenon arc lamp), according to (Rodrigues *et al*. 2017; Marques *et al*. 2019).

Data was recorded by a cooled CCD camera (Photometrics CoolSNAP). Before measurements, cell medium was replaced by warm standard buffer containing 119mM NaCl, 2mM Ca^2+^, 2mM MgCl2, 5mM KCl, 25mM HEPES, pH 7.4 (adjusted with NaOH). Cells were loaded with the Ca^2+^ sensitive dye Fura-2 AM (5μM, #47989, Life Technologies) for 45min before performing the intracellular calcium measurements. After 45min, Fura-2 AM was removed and replace by warm standard buffer (as described above) and experiments were performed at room temperature.

In order to evaluate the calcium response to the drugs, at 100s, an incubation with 1µM CBDV was performed for 700s. If the treatments required co-incubation with the antagonist 5’-IRTX, after the incubation with Fura-2 AM and 30min before performing the intracellular calcium measurements, cells were incubated with the antagonist at 300nM. At 800s, 2µM Ionomycin was added to record maximum response of cells (data not shown), as a positive control for cell response. Recordings ended after 900s.

Responses were recorded by a ratiometric method, in which image pairs were obtained every 5s by exciting the preparations at 340 and 380nm. Fura-2 AM has an absorbance at 340nm if bound to calcium, and at 380nm if not, while the emission wavelength is maintained at 510nm. The magnitude of the changes in the emission fluorescence of Fura-2 AM were taken as a measure of the changes in intracellular calcium concentration, as response to the drug stimulation.

### 2.10. Statistical analysis

Every independent experiment (*n*) corresponds to one independent SVZ neurosphere culture from one litter of C57BL/6J mice at P1-3. A minimum of 3 independent cultures was required to perform statistical analysis. Statistical analyses were performed using GraphPad Prism version 9.0.0 for Windows (GraphPad Software, Inc.), unless stated otherwise. Significance is reported as: ns: *p*>0.05; \**p*<0.05; \*\**p*<0.01; \*\*\**p*<0.001; \*\*\*\**p*<0.0001 when compared with control. § For comparisons of the co-incubation of drugs against the respective drug.

#### 2.10.1. For qRT-PCR experiments

SVZ-derived cells were grown in triplicates and each independent culture is considered n=1. Values were normalized to the control expression of GAPDH expression for each experiment. Data presented as Mean ± standard error of the mean (SEM) and the control was set to 1. Statistical significance was obtained using a Two-tailed Unpaired t-test.

#### 2.10.2. For the studies of cell viability, cell proliferation, neuronal/oligodendroglial differentiation, and pulse-chase experiments

All experiment measurements were performed at the border of SVZ neurospheres, where migrating cells form a pseudo-monolayer. Each condition was measured in three different coverslips. Percentages of immunoreactive cells were calculated from cell counts of five independent microscopic fields in each triplicated coverslip (representing n=1) with a 40x objective (≈100-200 cells per field). All experiments were analysed in a blind fashion and the obtained data was normalised to each corresponding control.

Normalisation of data was obtained by transformation using the Y=Y×K function in GraphPad Prism, with K=control group mean/100. Each individual experiment (*n*) was normalised to the respective control for that experiment. Normal distribution of data was tested using the Shapiro-Wilk normality test, for small sized samples (*n*<50) (Mishra *et al*. 2019). Data is presented as Mean ± SEM from the indicated number of independent cultures. For drug concentration-response curve experiments, statistical significance was obtained using a One-way ANOVA followed by Dunnett’s multiple comparisons *post-hoc* test. For the experiments with combination of drugs, statistical significance was obtained using a Two-way ANOVA followed by Bonferroni multiple comparisons *post-hoc* test. For the pulse-chase experiments, statistical significance was obtained using a Two-tailed Unpaired t-test. Outliers were identified and removed from the analysis by Mean ± (2 × standard deviation).

#### 2.10.3. For the neurosphere growth assay

Neurosphere diameter (DN, in µm) was measured using the Region of Interest (ROI) Manager tool from (Fiji Is Just) Image J version 2.3.0/1.53q for Windows OS (64-bit). The DN and the projected area (AN, in µm^2^) of each neurosphere were used as an indirect measure of cell proliferation. The size of the neurospheres was defined as an equivalent circle diameter, DN = 2(AN/π)^1/2^.

Neurospheres of DN < 30μm were excluded from the analysis because they were mainly single or paired cells. Data is presented as a violin plot with the median, 25% and 75% quartiles or as a bar chart representing the percentage of neurospheres binned according to size. The number of neurospheres considered for analysis in each condition (*n*) is from two independent cell cultures. Normal distribution of data was tested using the Kolmogorov–Smirnov test (*n*>50) (Mishra *et al*. 2019). Statistical significance was obtained using a Two-tailed Unpaired t-test. For the analysis of the comparison of proportions, statistical significance was obtained using the chi-square test using the MedCalc Software Ltd (MedCalc Software Ltd 2023), according to (Campbell 2007; Richardson 2011). Outliers were identified and removed from the analysis by Mean ± (2 × standard deviation).

#### 2.10.4. For calcium imaging experiments

Images were collected and analysed using MetaFluor Fluorescence Ratio Imaging Software (Molecular Devices). Regions of interest were acquired by delineating the profile of the cells and averaging the fluorescence intensity inside the delineated area. Statistical significance was obtained after peak determination by the analysis of the area under the curve. Peak amplitudes were calculated by subtracting the baseline level to the maximum peak intensity. Experiments were performed at least in triplicate, except stated otherwise. The number of responsive cells is designated by *n*. A cell was considered responsive when its maximum recorded response was greater than the average of the responses for all cells for each condition. Data is expressed as Mean ± SEM or as a bar chart representing the proportion of responsive vs non-responsive cells. Statistical significance was obtained using a Two-way ANOVA followed by Bonferroni multiple comparisons *post-hoc* test. For the analysis of the comparison of proportions, statistical significance was obtained using the chi-square test using the MedCalc Software Ltd (MedCalc Software Ltd 2023), according to (Campbell 2007; Richardson 2011).

## 3. Results

### 3.1. TRPV1 expression is increased by CBDV

To evaluate the expression of TRPV1 in our culture model, SVZ neurospheres were incubated for 7 days *in vitro* (DIV7) under differentiative conditions, in the presence or absence of CBDV, and further processed for qRT-PCR. We observed that not only TRPV1 is expressed in SVZ-derived cells, but also its expression is increased in the presence of CBDV, when compared to the control condition (t(6)=5.487, *p*=0.0015; Ctrl: 1.0-fold; CBDV 1µM: 2.457±0.363-fold; n=3-5; \*\*\**p*<0.001) (Fig. S1).

### 3.2. CBDV promotes cell viability via a TRPV1-dependent mechanism of action

The effect of CBDV in cell viability was studied using a drug concentration-response curve in SVZ-derived cells. For that, SVZ neurospheres were incubated for 2 days *in vitro* (DIV2) under differentiative conditions with increasing concentrations of CBDV (100nM - 1µM) and, 30 minutes before fixing, cells were incubated with Propidium Iodide (PI) to label late apoptosis or necrosis (Lecoeur 2002) (Fig. 1A, 1D).

**Fig. 1.**
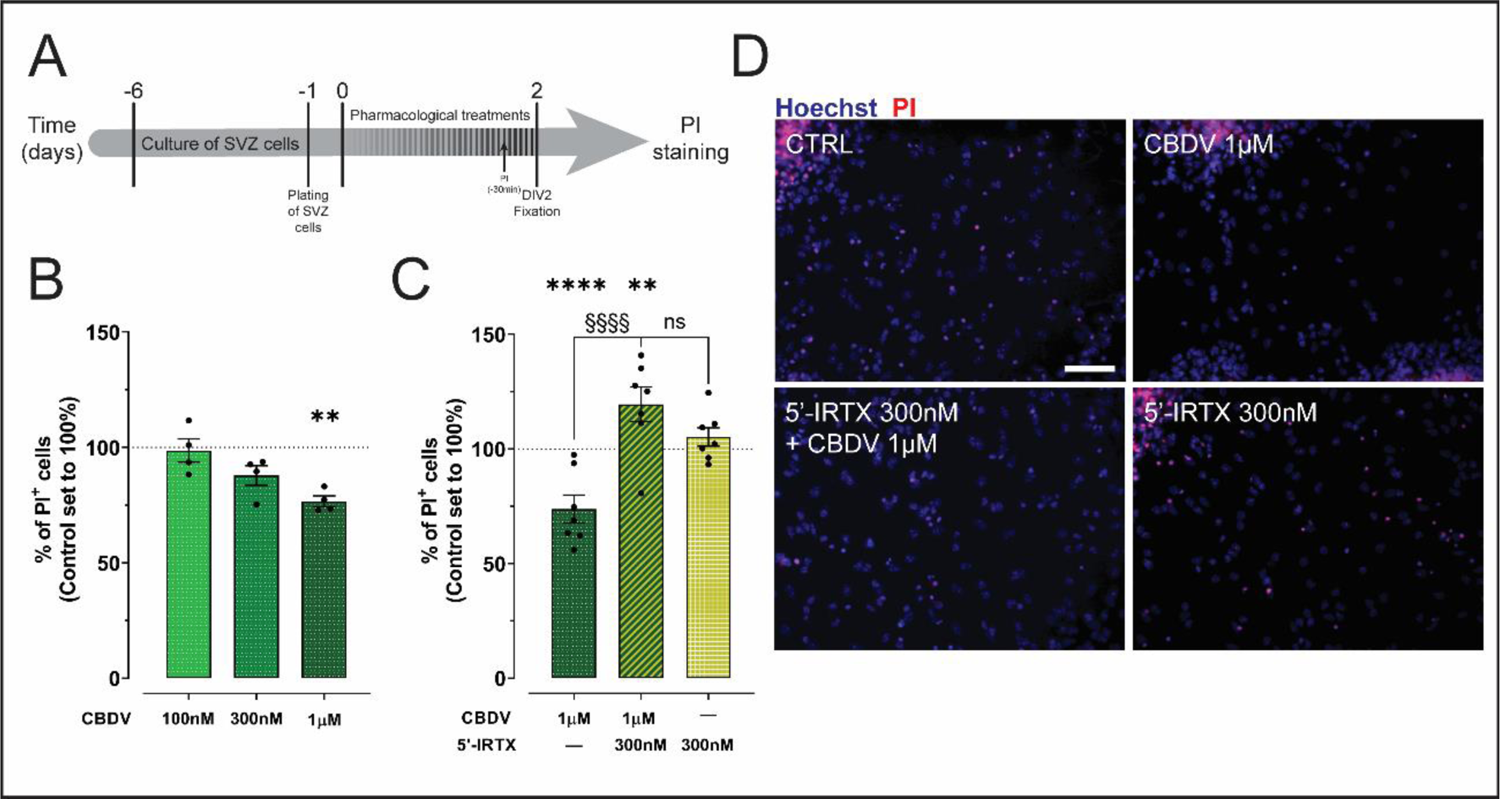
Cannabidivarin promotes cell viability under differentiative conditions. **(A)** Schematic representation of the protocol used to evaluate cell viability. **(B)** Bar graphs depict the percentage of PI^+^ cells treated with CBDV for DIV2. Values were normalized to the control mean for each experiment. Data presented as Mean ± SEM and the control was set to 100% (corresponding to 15.72% PI^+^ cells). n=4; **p<0.01. One-way ANOVA using Dunnett’s multiple comparison *post-hoc* test. **(C)** Bar graphs depict the percentage of PI^+^ cells co-treated with 5’-iRTX and CBDV for DIV2. Values were normalized to the control mean for each experiment. Data presented as Mean ± SEM and the control was set to 100%. n=7; ns: p>0.05; **p<0.01; ****p<0.0001; §§§§p<0.0001. Two-way ANOVA followed by Bonferroni multiple comparisons *post-hoc* test. **(D)** Representative fluorescent images of cells positive for PI (in red) and Hoechst 33342 staining (blue nuclei) at DIV2. Scale bar=50μm.

At DIV2, CBDV-treated cells, at the highest concentration, showed a reduction in the percentage of PI^+^ cells when compared to the control condition, therefore promoting cell survival (F(3,12)=9, *p*=0.0016; Ctrl: 100±0.0112%; CBDV 1µM: 76.63±2.379%; n=4; \*\**p*<0.01) (Fig. 1B, 1D). To evaluate whether the effects of CBDV upon cell viability were TRPV1-dependent, cells were incubated with the TRPV1 antagonist 5’-iodoresinferotoxin (5’-IRTX) at 300nM for 30 minutes, before being co-incubated with the highest concentration of CBDV (1µM). For CBDV-induced cell survival, the decrease in cell death was blocked in the presence of the antagonist, revealing that CBDV-induced cell survival is TRPV1-dependent (CBDV×5’-IRTX: F(3,50)=23.19, *p*<0.0001; CBDV 1µM: 73.76±6.068%; 5’-IRTX 300nM + CBDV 1µM: 119.4±7.555%; n=7; §§§§*p*<0.0001) (Fig. 1C, 1D). 5’-IRTX alone, in turn, had no effect on cell viability when compared to the control condition (Fig. 1C, 1D).

Taken together, this data demonstrates that CBDV is able to regulate the viability of SVZ-derived cells via a TRPV1-dependent mechanism of action.

### 3.3. CBDV promotes cell proliferation in a TRPV1-dependent mechanism of action

To study the effect of CBDV in cell proliferation, SVZ neurospheres were incubated with increasing concentrations of CBDV (100nM - 1µM) for DIV2 under differentiative conditions and, 4h before fixing, cells were incubated with 5-bromo-2’-deoxyuridine (BrdU) to label proliferating cells (Fig. 2A, 2E).

**Fig. 2.**
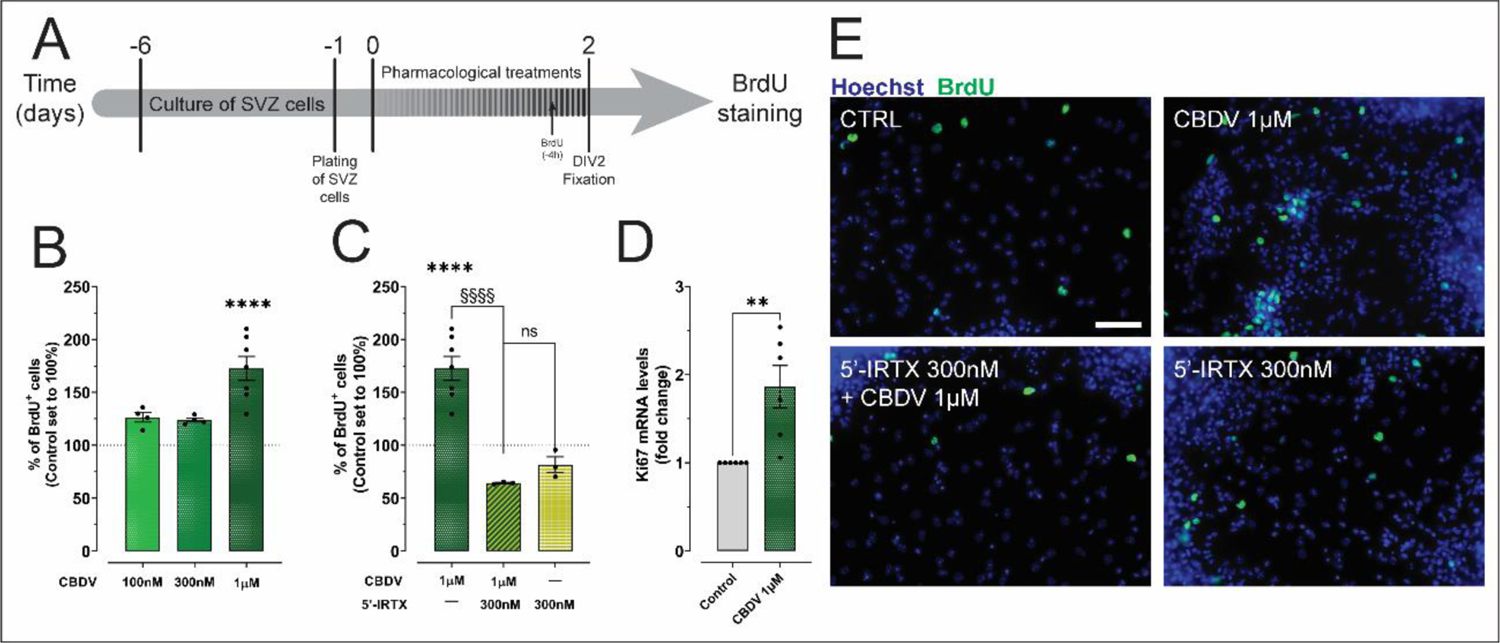
Cell proliferation is increased with Cannabidivarin-treatment under differentiative conditions. **(A)** Schematic representation of the protocol used to evaluate cell proliferation. **(B)** Bar graphs depict the percentage of BrdU^+^ cells treated with CBDV for DIV2. Values were normalized to the control mean for each experiment. Data presented as Mean ± SEM and the control was set to 100% (corresponding to 6.39% BrdU^+^ cells). n=4-7; ****p<0.0001. One-way ANOVA using Dunnett’s multiple comparison *post-hoc* test. **(C)** Bar graphs depict the percentage of BrdU^+^ cells co-treated with 5’-iRTX and CBDV for DIV2. Values were normalized to the control mean for each experiment. Data presented as Mean ± SEM and the control was set to 100%. n=3-7; ns: p>0.05; ****p<0.0001; §§§§p<0.0001. Two-way ANOVA followed by Bonferroni multiple comparisons *post-hoc* test. **(D)** Bar graph depicts the levels of Ki67 mRNA expression. Values were normalized to the control expression of GAPDH expression for each experiment. Data presented as Mean ± SEM and the control was set to 1. n=6; **p<0.01. Unpaired t test. **(E)** Representative fluorescent images of cells positive for BrdU (in green) and Hoechst 33342 staining (blue nuclei) at DIV2. Scale bar=50μm.

CBDV-treated cells, at the highest concentration, showed an increase in the percentage of BrdU^+^ cells when compared to the control condition (F(3,17)=8.937, *p*=0.0009; Ctrl: 100±0.00182%; CBDV 1µM: 172.6±11.360%; n=4-7; \*\*\*\**p*<0.0001) (Fig. 2B, 2E). Importantly, when the antagonist 5’-IRTX was co-incubated with CBDV, the increase in CBDV-induced cell proliferation was blocked (CBDV×5’-IRTX: F(3,41)=56.25, *p*<0.0001; CBDV 1µM: 172.6.11±11.360%; 5’-IRTX 300nM + CBDV 1µM: 64.36±0.6442%; n=3-7; §§§§*p*<0.0001) (Fig. 2C, 2E). Incubation with the antagonist 5’-IRTX alone had no effect on NSPC proliferation (Fig. 2C, 2E).

To test whether the CBDV-induced proliferative properties in NSPCs were long-lasting and to further assess cell proliferation by a different technique at a different time-point, the expression of Ki67 mRNA, a nuclear protein associated with cellular proliferation, was evaluated in SVZ-derived cells at DIV7. Corroborating the BrdU assays at DIV2, a significant increase in Ki67 expression levels was observed in CBDV-treated cells (t(10)=3.552, *p*=0.0053; Ctrl: 1.0-fold; CBDV 1µM: 1.862±0.242-fold; n=6; \*\**p*<0.01) (Fig. 2D).

Furthermore, cell proliferation was also evaluated under proliferative conditions through the assessment of neurosphere size (Fig. 3A, 3E). In agreement with the results obtained under differentiative conditions, CBDV significantly increased the size of neurospheres, both in terms of diameter (DN, Fig. 3B) and area (AN, Fig. 3C), when compared to the control condition (DN: t(3829)=7.658, *p*<0.0001; Ctrl: 94.36 (70.81 to 124.0)µm, n=2110; CBDV 1µM: 100.5 (75.49 to 140.1)µm, n=1721; \*\*\*\**p*<0.0001; AN: t(3900)=8.629, *p*<0.0001; Ctrl: 7143 (3987 to 12526)µm^2^, n=2143; CBDV 1µM: 8177 (4535 to 16195)µm^2^, n=1759; \*\*\*\**p*<0.0001). A more in-depth analysis of the DN revealed that CBDV-treated neurospheres generate more neurospheres with a DN > 300µm than control condition (*Χ^2^* (1, *N=*3969)=20.761, *p*<0.0001; Ctrl: 1.37%, n=2177; CBDV 1µM: 3.58%, n=1792) (Fig. 3D, 3E; Table S1).

**Fig. 3.**
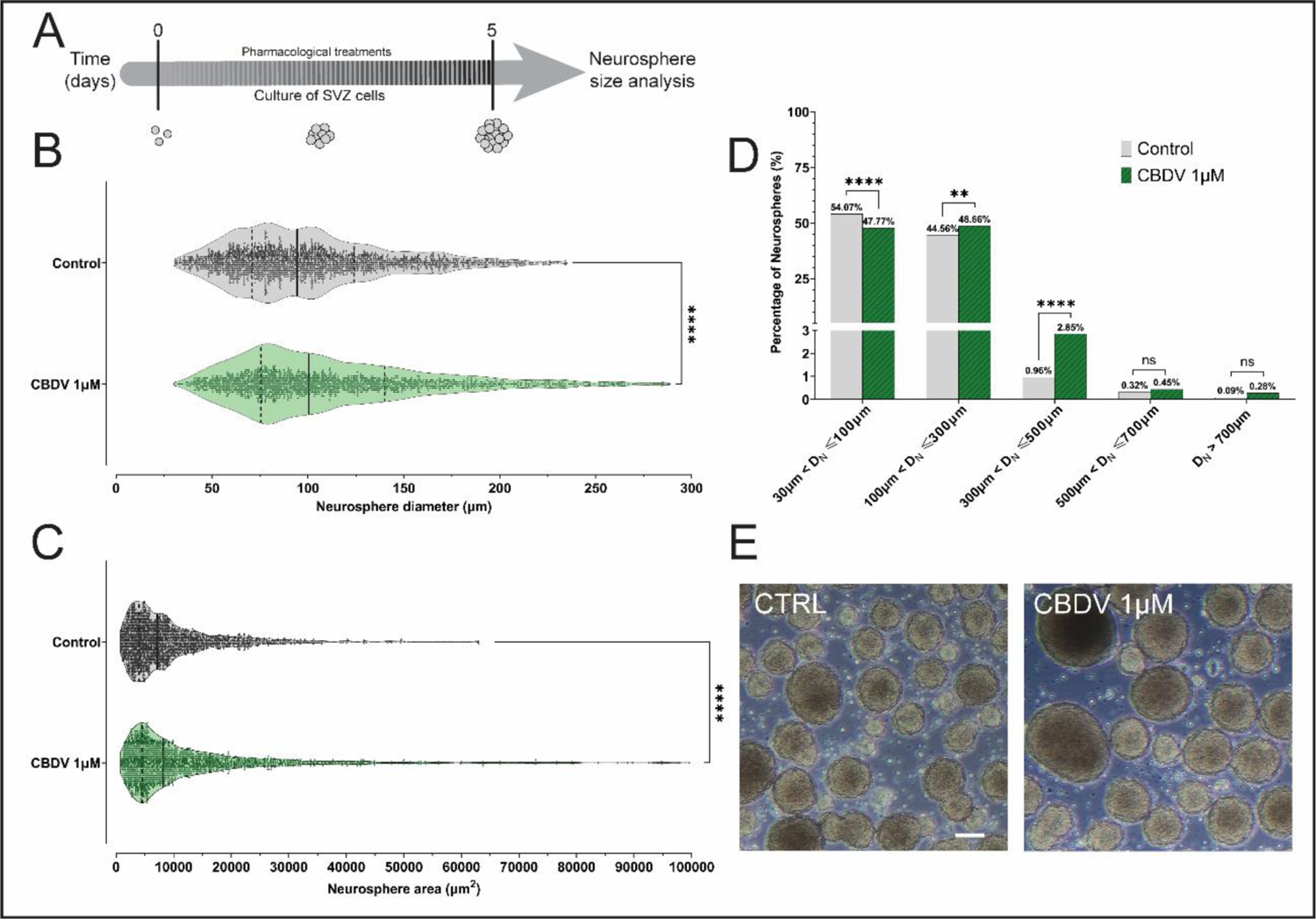
Neurosphere size is increased with Cannabidivarin-treatment under proliferative conditions. **(A)** Schematic representation of the protocol used to evaluate neurosphere size. **(B)** Violin plot representing the variation of neurosphere diameter (DN). Data presented as median, 25% and 75% quartiles. The total number of neurospheres analysed (n) is for Ctrl: 2110, CBDV: 1721, from two independent cultures, ****p<0.0001. Unpaired t test. **(C)** Violin plot representing the variation of neurosphere area (AN). Data presented as median, 25% and 75% quartiles. The total number of neurospheres analysed (n) is for Ctrl: 2143, CBDV: 1759, for two independent cultures, ****p<0.0001. Unpaired t test. **(D)** Bar graph representing the percentages of neurospheres distributed according to the size-binning categories, ns: p>0.05; **p<0.01; ****p<0.0001. Chi-square test. **(E)** Representative phase contrast images of neurospheres after 5 days in culture. Scale bar=100μm.

Thus, these results demonstrate that CBDV promotes cell proliferation in both proliferative and differentiative conditions in a TRPV1-dependent mechanism of action.

### 3.4. CBDV promotes neuronal differentiation in a TRPV1-dependent mechanism of action

Neuronal differentiation was evaluated in SVZ neurospheres treated with increasing concentrations of CDBV (100nM - 1µM), for DIV2 or DIV7, under differentiative conditions. For that, immunocytochemistry (ICC) assays against NeuN, a marker for mature neurons, were performed (Fig. 4A, 4F).

**Fig. 4.**
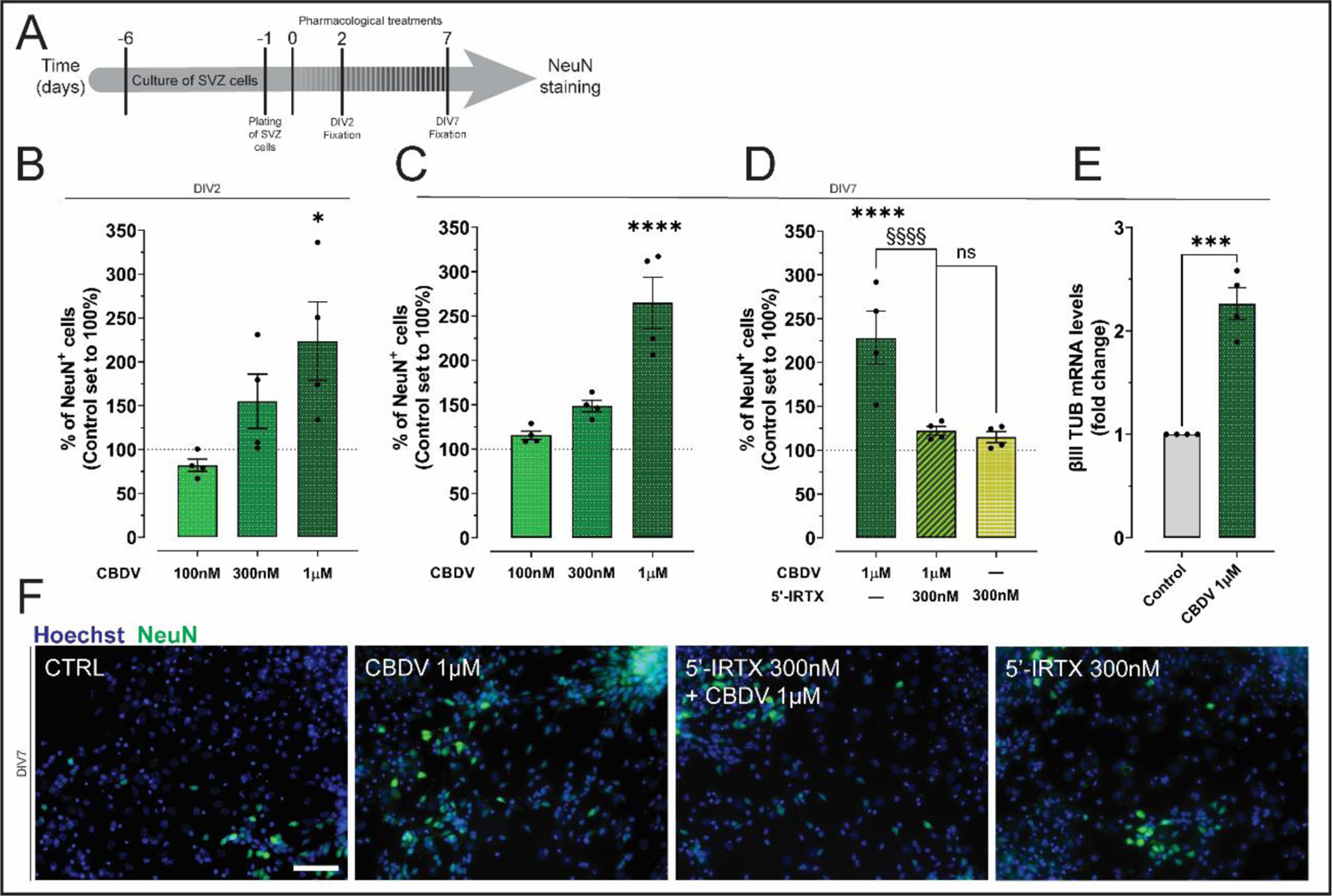
Neuronal differentiation is increased by Cannabidivarin treatment. **(A)** Schematic representation of the protocol used to evaluate neuronal differentiation. **(B)** Bar graphs depict the percentage of NeuN^+^ cells treated with CBDV for DIV2. Values were normalized to the control mean for each experiment. Data presented as Mean ± SEM and the control was set to 100% (corresponding to 3.468% NeuN^+^ cells). n=4; *p<0.05. One-way ANOVA using Dunnett’s multiple comparison *post-hoc* test. **(C)** Bar graphs depict the percentage of NeuN^+^ cells treated with CBDV for DIV7. Values were normalized to the control mean for each experiment. Data presented as Mean ± SEM and the control was set to 100% (corresponding to 11.143% NeuN^+^ cells). n=4; *p<0.05; ****p<0.000. One-way ANOVA using Dunnett’s multiple comparison *post-hoc* test. **(D)** Bar graphs depict the percentage of NeuN^+^ cells co-treated with 5’-IRTX and CBDV for DIV7. Data presented as Mean ± SEM and the control was set to 100%. n=4; ns: p>0.05; ****p<0.0001; §§§§p<0.0001. Two-way ANOVA followed by Bonferroni multiple comparisons *post-hoc* test. **(E)** Bar graph depicts the levels of βIII-tubulin mRNA expression. Values were normalized to the control expression of GAPDH expression for each experiment. Data presented as Mean ± SEM and the control was set to 1. n=4; ***p<0.001. Unpaired t test. **(F)** Representative fluorescent images of cells positive for NeuN (in green) and Hoechst 33342 staining (blue nuclei) at DIV7. Scale bar=50μm.

Our results showed that the highest concentration of CBDV increased the percentage of NeuN^+^ cells at DIV2, when compared to the control condition (F(3,12)=5.422, *p*=0.0137; Ctrl: 100±0.00583%; CBDV 1µM: 223.7±44.690%; n=4; \**p*<0.05) (Fig. 4B).

Importantly, at DIV7, the effect of the highest concentration of CBDV in neuronal differentiation was still detected (F(3,12)=24.67, *p*<0.0001; Ctrl: 100±0.00139%; CBDV 1µM: 264.8±28.930%; n=4; \*\*\*\**p*<0.0001) (Fig. 4C, 4F). Furthermore, at this time point, blocking TRPV1 with 5’-IRTX alone had no significant differences in the percentage of neurons when compared to the control condition. Notwithstanding, CBDV-induced neuronal differentiation was blocked with TRPV1 antagonist (CBDV×5’-IRTX: F(3,26)=62.16, *p*<0.0001; CBDV 1µM: 228.1±30.480%; 5’-IRTX 300nM + CBDV 1µM: 122.1±4.984%; n=4; §§§§*p*<0.001) (Fig. 4D, 4F).

To further understand the effects of CBDV and Capsaicin on the degree of neuronal maturation, the mRNA expression levels of βIII-tubulin, a marker for immature neurons, was evaluated in SVZ neurospheres at DIV7 under differentiative conditions. In fact, in CBDV-treated cells, the mRNA expression of βIII-tubulin was found significantly increased (t(6)=8.177, *p*=0.0002; Ctrl: 1.0-fold; CBDV 1µM: 2.263±0.154-fold; n=4; \*\*\**p*<0.001) (Fig. 4E).

These results highlight the importance of TRPV1 in neuronal differentiation process, namely on its impact in regulating the early stages of NSPC pool and maturation of newborn neurons.

### 3.5. CBDV-induced neuronal differentiation is a result of cell cycle exit of NSPCs

To further clarify whether the CBDV-induced increase in neuronal differentiation relies on their effects in controlling proliferating and/or cell cycle exit of NSPCs, neurospheres, under differentiative conditions, were subjected to CBDV treatment and exposed to a pulse of BrdU for the first 24h. The drugs, however, remained in culture until DIV7, the time at which cells were co-labelled against BrdU and NeuN (Fig. 5A, 5F).

**Fig. 5.**
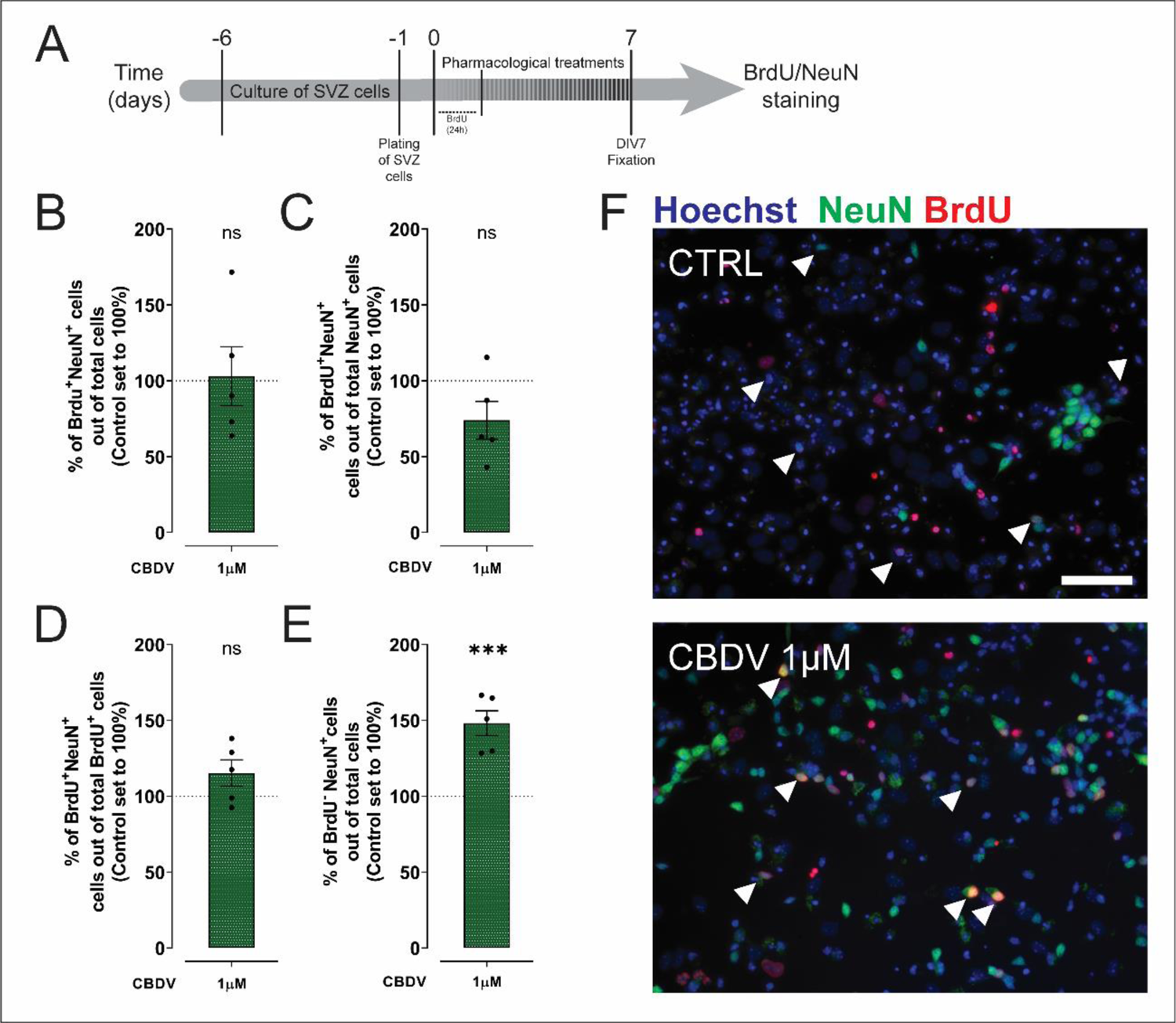
CBDV promotes NSPC exit of the cell cycle and differentiation into neurons. **(A)** Schematic representation of the protocol used to evaluate differentiation of neurons from proliferative NSPCs. **(B)** Bar graphs depict the percentage of BrdU^+^NeuN^+^ cells out of total cells. **(C)** Bar graphs depict the percentage of BrdU^+^NeuN^+^ cells out of NeuN^+^. **(D)** Bar graphs depict the percentage of BrdU^+^NeuN^+^ cells out of BrdU^+^. **(E)** Bar graphs depict the percentage of BrdU^‒^NeuN^+^ out of total cells. **(B-E)** Values were normalized to the control mean for each experiment. Data presented as Mean ± SEM and the control was set to 100% (corresponding to **(B)** 2.490% BrdU^+^NeuN^+^ cells out of total cells; **(C)** 15.731% BrdU^+^NeuN^+^ cells out of NeuN^+^; **(D)** 38.416% BrdU^+^NeuN^+^ cells out of BrdU^+^; **(E)** 13.746% BrdU^‒^NeuN^+^ out of total cells); n=5; ns: p>0.05; ***p<0.001. Unpaired t test. **(F)** Representative fluorescent images of cells positive for NeuN (in green), BrdU (in red) and Hoechst 33342 staining (blue nuclei) at DIV 7. Arrows represent double-positive cells for BrdU/NeuN. Scale bar=50μm.

Surprisingly no significant changes in the percentage of BrdU^+^NeuN^+^ cells (out of total cells) were observed in CBDV condition (t(8)=0.1553, *p*=0.8804) (Fig. 5B, 5F). When looking at the percentage of neurons that differentiated from proliferating cells (BrdU^+^NeuN^+^ cells (out of NeuN^+^)) (Fig. 5C, 5F), again no differences were observed (t(8)=2.080, *p*=0.0711), suggesting that the differentiated BrdU^+^NeuN^+^ neurons represent only a small fraction of the total population of neurons. Concomitantly, when looking at the percentage of proliferating cells that differentiated into neurons (BrdU^+^NeuN^+^ cells (out of BrdU^+^)) (Fig. 5D, 5F) no differences were observed when comparing to the control condition (t(8)=1.760, *p*=0.1164), suggesting that CBDV-induced proliferative cells are not responsible for the increased percentage of neurons observed after this treatment. However, an increase in the percentage of BrdU^‒^NeuN^+^ cells (out of total cells) by CBDV treatment was observed (t(8)=5.865, *p*=0.0004; Ctrl: 100±0.01139%; CBDV 1µM: 148.2±8.213%; n=5; \*\*\**p*<0.001), revealing that CBDV induced the arrest and exit of cell cycle in NSPCs and stimulates their differentiation into neurons (Fig. 5E, 5F).

In conclusion, our BrdU/NeuN experiments clearly show that CBDV-increased NeuN^+^ cells result from NSPCs that exit the cell cycle.

### 3.6. CBDV inhibits oligodendrocyte differentiation and maturation

Since cannabinoids have been shown to regulate oligodendrocyte differentiation in the SVZ (Arévalo-Martín *et al*. 2007), we aimed at understanding if CBDV would share this effect and if TRPV1 was involved. To do so, neurospheres, under differentiative conditions, were treated with CBDV for DIV2 and DIV7, and an ICC against Neuron/glia antigen 2 (NG2), a marker for oligodendrocyte progenitor cells (OPC), was performed (Fig. 6A, 6F). Notably, CBDV did not induced significant changes in the percentage of NG2^+^ cells at DIV2 when compared to the control condition (F(3,12)=0.1647, *p*=0.9181) (Fig. 6B). However, at DIV7, the highest concentration of CBDV (F(3,12)=12.12, *p*=0.0006; Ctrl: 100±0.0107%; CBDV 1µM: 72.61±3.506%; n=4; \*\**p*<0.01) significantly reduced the percentage of OPCs (Fig. 6C, 6F). Therefore, given the reduction in the percentage of OPCs, we hypothesized that CBDV could be promoting of oligodendrocyte maturation. Thus, an ICC was performed in the same conditions at DIV7, but this time against Myelin Basic Protein (MBP), a marker for mature myelinating oligodendrocytes (Fig. 6A, 6G). Surprisingly, the highest concentration of CBDV diminished the percentage of MBP^+^ cells (F(3,12)=10.80, *p*=0.0010; Ctrl: 100±0.00372%; CBDV 1µM: 44.65±3.506%; n=4; \*\**p*<0.01) (Fig. 6D, 6G).

**Fig. 6.**
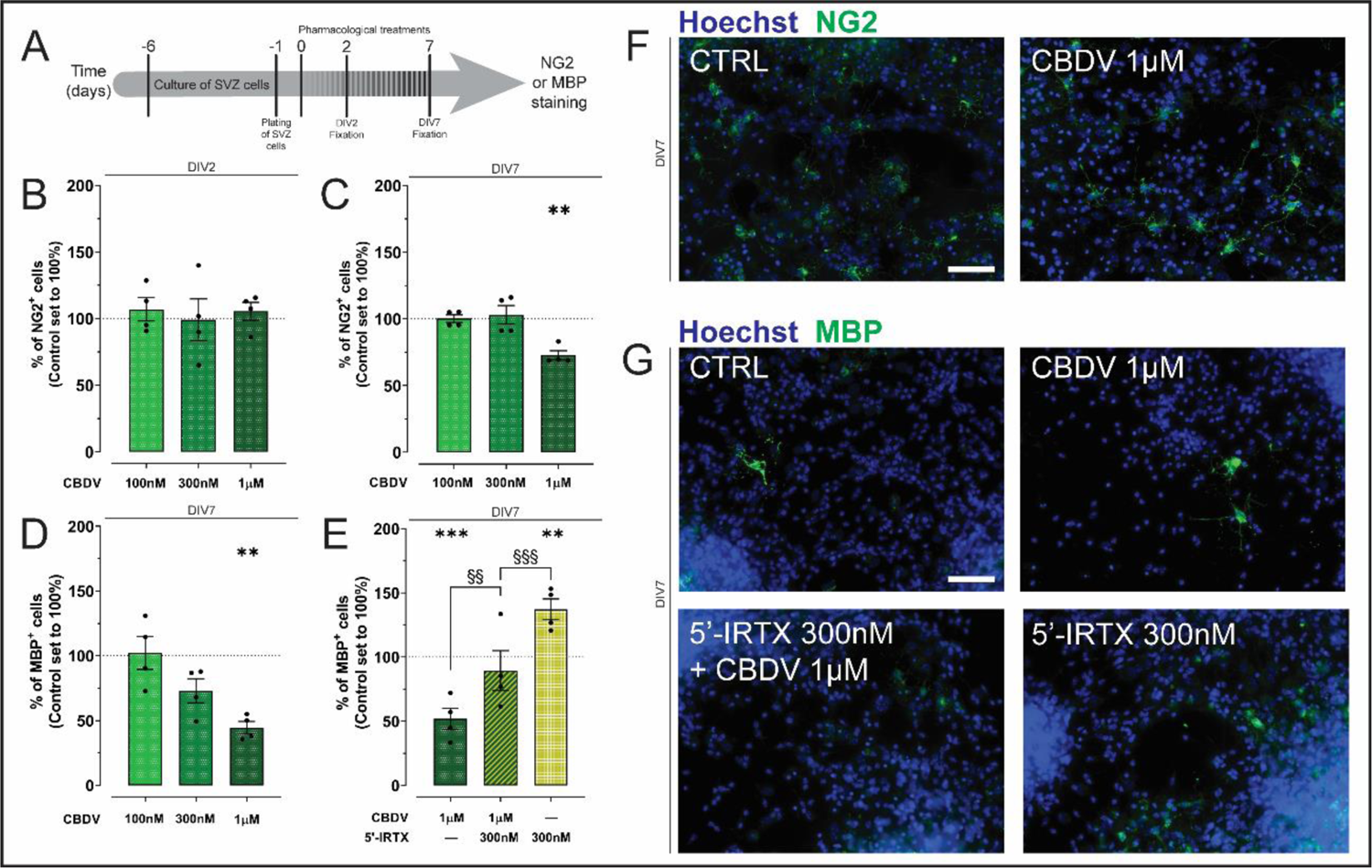
Oligodendroglial differentiation and maturation is inhibited by Cannabidivarin treatment. **(A)** Schematic representation of the protocol used to evaluate oligodendrocyte differentiation. **(B)** Bar graphs depict the percentage of NG2^+^ cells treated with CBDV for DIV2. Values were normalized to the control mean for each experiment. Data presented as Mean ± SEM and the control was set to 100% (corresponding to 13.24% NG2^+^ cells). n=4; ns: p>0.05. One-way ANOVA using Dunnett’s multiple comparison *post-hoc* test. **(C)** Bar graphs depict the percentage of NG2^+^ cells treated with CBDV for DIV7. Data presented as Mean ± SEM and the control was set to 100% (corresponding to 6.943% NG2^+^ cells). n=4; **p<0.01. One-way ANOVA using Dunnett’s multiple comparison *post-hoc* test. **(D)** Bar graphs depict the percentage of MBP^+^ cells treated with CBDV for DIV7. Data presented as Mean ± SEM and the control was set to 100% (corresponding to 1.395% of MBP^+^ cells). n=4; **p<0.01. One-way ANOVA using Dunnett’s multiple comparison *post-hoc* test. **(E)** Bar graphs depict the percentage of MBP^+^ cells co-treated with 5’-IRTX and CBDV for DIV7. Data presented as Mean ± SEM and the control was set to 100%. n=4; ns: p>0.05; **p<0.01; ***p<0.001; §§p<0.01; §§§p<0. 001. Two-way ANOVA followed by Bonferroni multiple comparisons *post-hoc* test. **(F)** Representative fluorescent images of cells positive for NG2 (in green) and Hoechst 33342 staining (blue nuclei) at DIV7. **(G)** Representative fluorescent images of cells positive for MBP (in green) and Hoechst 33342 staining (blue nuclei) at DIV7. Scale bars=50μm.

In addition, and strikingly, blocking TRPV1 with the antagonist 5’-IRTX was able to increase the percentage of MBP^+^ cells, when compared to the control condition (F(3,12)=13.25, *p*=0.0004; Ctrl: 100± 0.00134%; 5’-IRTX 300nM: 137.3±7.918%; n=4; \*\**p*<0.01) (Fig. 6E, 6G), suggesting that TRPV1 inhibition *per se* promotes MBP differentiation and maturation. As such, when co-incubating 5’-IRTX with CBDV, an increase in the percentage of myelinating oligodendrocytes was detected when comparing with CBDV alone (CBDV×5’-IRTX: F(3,28)=21.79, *p*<0.0001; CBDV 1µM: 51.81±8.299%; 5’-IRTX 300nM + CBDV 1µM: 89.34±15.55%; n=4; §§*p*<0.01) (Fig. 6E, 6G).

These results reveal a potential for TRPV1 in the modulation of oligodendrocyte differentiation and maturation.

### 3.7. CBDV-responsive cells display a two-phase calcium influx profile

Since CBDV acts via TRPV1, a sodium-calcium channel, and that intracellular calcium levels are known regulators of NSPCs fate and neuronal maturation, single cell calcium imaging was performed to evaluate the functional response of SVZ-derived cells at DIV7 exposed, or not (Movie A1) to CBDV. Cells were also exposed to a prior incubation/non-incubation with the TRPV1 antagonist, 5’-ITRX, to further clarify the role of TRPV1 activation in response to these drugs (Fig. 7A).

**Fig. 7.**
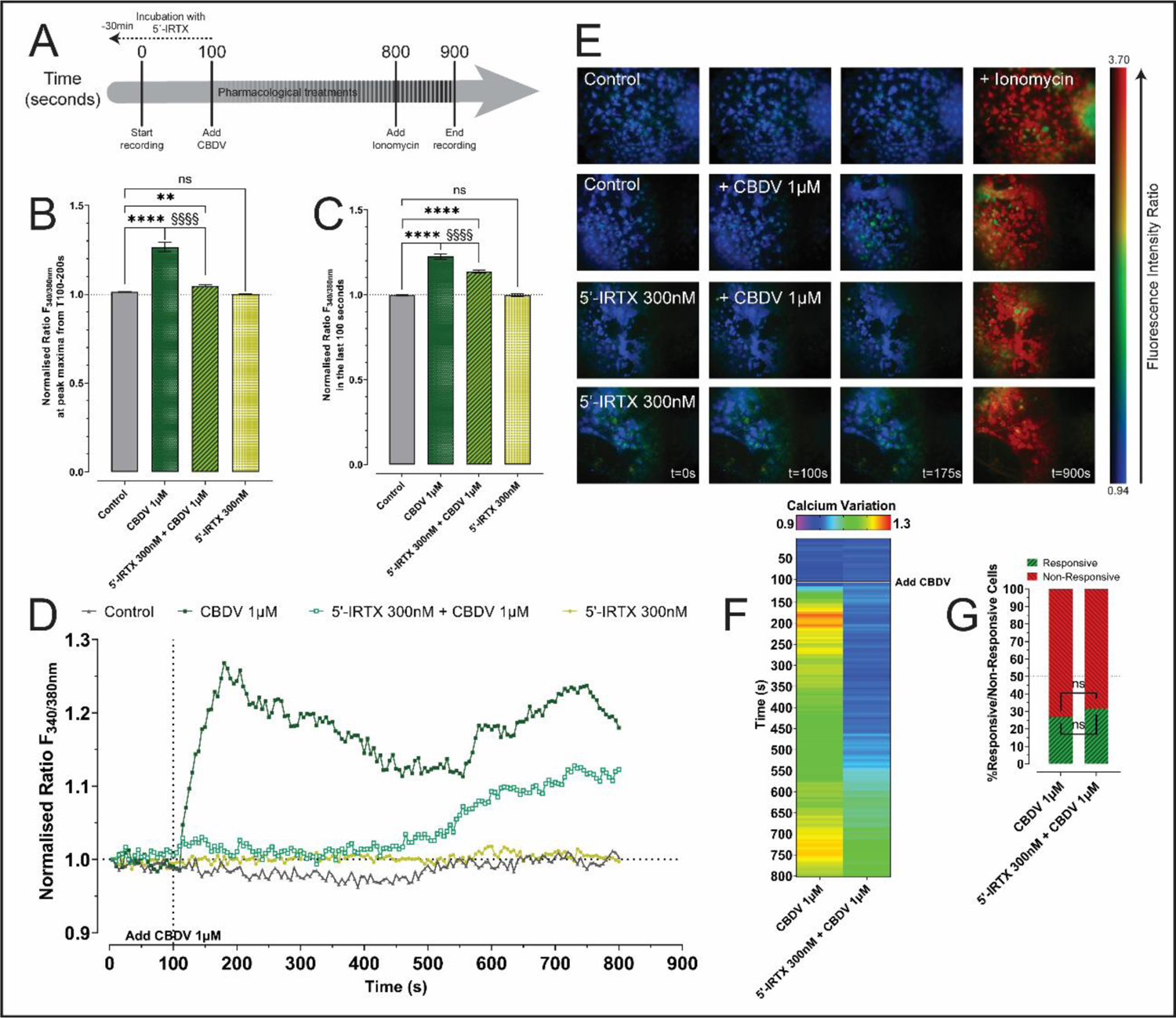
CBDV-induced calcium influx profile is dependent of TRPV1 activation in SVZ-derived cells. **(A)** Schematic representation of the protocol used to evaluate calcium influx. **(B)** Ratio of calcium influx at peak maxima from T=100-200 seconds, after CBDV exposure. **(C)** Ratio of calcium influx at peak maxima in the last 100s of experiment, after CBDV exposure. **(B)(C)** The responding cells were analysed in each well (120-365 cells) and normalized to the first point of the baseline for each data set. n=3; ns: p>0.05; **p<0.01; ****p<0.01; §§§§p<0.0001. Two-way ANOVA followed by Bonferroni multiple comparisons post-hoc test at peak maxima. **(D)** Representative scatter plot that depicts a single-cell variation of calcium influx throughout the time (in seconds) in response to 1µM CBDV. **(E)** Representative fluorescent intensity ratio of calcium influx (colour code, blue for a low calcium level, green for intermediate, yellow for medium high, and red for high). **(F)** Representative matrix of single-cell calcium response to a pulse of 1µM CBDV. Each row represents time-points, and the columns represent the mean values of calcium influx of most responsive cells in each condition. The heatmap graph represents a colour-code variation (purple/blue for a low calcium level, green for intermediate, yellow for medium high, and red for high) of the mean values of calcium influx of the most responsive cells from a total of 973 cells analysed in these conditions. **(G)** Stacked bar graphs representing the percentage of responsive versus non-responsive cells influx upon a pulse of 1µM CBDV. A cell was considered responsive when its maximum recorded response was greater than the average of the responses for all cells for each condition. ns: p>0.05; Chi-square test.

Surprisingly, incubation with 1µM CBDV at 100 seconds elicited two types of calcium influx responses in SVZ-derived cells. One short initial response (≈85 seconds after the CBDV was added to cells), that corresponds to TRPV1 activation, which is blocked by the antagonist 5’-IRTX, and a second long-lasting response (≈550 seconds after CBDV was added to cells). Specifically, in the first response, CBDV elicited a moderate increase in the calcium influx, with a peak, followed by a subtle decline (F(3,976)=200.5, *p*<0.0001; Ctrl peak ratio: 1.014±0.00173, n=365; CBDV 1µM peak ratio: 1.266±0.0260, n=120 responsive cells; \*\*\*\**p*<0.0001). In contrast, in cells co-exposed to CBDV and 5’-IRTX, the initial calcium influx was blocked (CBDV×5’-IRTX: F(3,612)=145.4, *p*<0.0001; CBDV 1µM peak ratio: 1.266±0.0260, n=120; 5’-IRTX 300nM + CBDV 1µM peak ratio: 1.049±0.00598, n=166 responsive cells; §§§§*p*<0.0001). Unexpectedly, an increase in calcium influx, starting at around 500 seconds, was also observed, peaking at ≈650s, after CBDV was added, lasting until the end of the experiment. Of note, when compared to the control condition, the co-exposure with the antagonist, resulted in a slight increase of calcium influx at the peak (CBDV×5’-IRTX: F(3,612)=145.4, *p*<0.0001; Ctrl peak ratio: 1.014±0.00173, n=365; 5’-IRTX 300nM + CBDV 1µM peak ratio: 1.049±0.00598, n=166 responsive cells; \*\**p*<0.01) (Fig. 7B, 7D, 7E, 7F, Movie A2, Movie A4). No significant differences in the calcium profile were detected when comparing the calcium influx of 5’-IRTX exposed cells with the control condition at the peak (Movie A3). Similarly, in the last 100 seconds of responses, no differences in the calcium influx were observed when comparing control with 5’-IRTX alone (Movie A3). Moreover, the calcium influx induced by CBDV was higher than the influx induced by the co-exposure with the antagonist in the last 100 seconds of responses (CBDV×5’-IRTX: F(3,377)=115.7, *p*<0.0001; CBDV 1µM ratio: 1.224±0.0160, n=120; 5’-IRTX 300nM + CBDV 1µM: 1.136±0.00908, n=166 responsive cells; §§§§p<0.0001). Furthermore, the influx induced by both CBDV or by co-exposure with the antagonist did not reach levels close to the control baseline in the last 100 seconds (F(3,741)=178.9, *p*<0.0001; CBDV×5’-IRTX: F(3,377)=115.7, *p*<0.0001; Ctrl ratio: 0.9964±0.00368, n=365; CBDV 1µM ratio: 1.224±0.0160, n=120; ****p<0.0001; 5’-IRTX 300nM + CBDV 1µM ratio: 1.136±0.00908, n=166 responsive cells; ****p<0.0001) (Fig. 7C, 7D, 7E, 7F, Movie A2, Movie A4).

Interestingly, out of all the analysed cells, no differences were found between the percentage of responsive and non-responsive cells. In more detail, when comparing cells that responded to CBDV to those that were co-incubated with the antagonist, the proportion of responsive cells was equivalent (*Χ^2^* (1, *N=*973)=0.839, *p*=0.3596; %Responsive CBDV 1µM: 26.85%; %Responsive 5’-IRTX 300nM + CBDV 1µM: 31.56%). Concomitantly, the proportion on non-responsive cells also presented no differences (*Χ^2^* (1, *N=*973)=2.580, *p*=0.1082; %Non-Responsive CBDV 1µM: 73.15%; %Non-Responsive 5’-IRTX 300nM + CBDV 1µM: 68.44%) (Fig. 7G).

In conclusion, CBDV was able to induce two types of response. One, in the first 100s after CBDV exposure, that was blocked by 5’-IRTX, and another that was not blocked by the antagonist in the last 100s of the experimental protocol.

## 4. Discussion

In this study we describe the proneurogenic effects of CBDV, *in vitro*, as a direct effect of TRPV1 activation. CBDV positively regulates cell survival, cell proliferation and neuronal differentiation in a TRPV1-dependent mechanism of action. Furthermore, inhibition of oligodendrogenesis by CBDV was also observed, which was blocked with TRPV1 antagonist.

Although the affinity of CBDV for TRPV1 is yet uncertain, since the KiTRPV1 for CBDV has not been yet determined, the concentrations of CBDV used in this study were based on previous works made with other endo-, phytho- and synthetic cannabinoids (Xapelli *et al*. 2013; Stanslowsky *et al*. 2017; Compagnucci *et al*. 2013; Rodrigues *et al*. 2017). In an elegant study by Petrocellis and colleagues (De Petrocellis *et al*. 2011), they determined the EC50 for CBDV for several receptors, namely TRPV1, TRPV2 and TRPA1. Their study has served as the base of several others that further clarified the pharmacology of this drug. Despite that CBDV was reported to have a EC50 for TRPA1 (0.42±0.01µM) lower than for TRPV1 (3.6±0.7µM), it is possible that these results might be influenced by the methodology applied to determine the EC50 (MarÉchal 2011). In their study, HEK-293 cells were transfected using recombinant rat TRPA1, rat TRPV2 and human TRPV1. While, not only these cells do not physiologically express these receptors (Costanzo *et al*. 2020; Starkus *et al*. 2019), the constructs used were from different species, thus the calcium changes, used to determine the EC50, might not totally translate into physiological responses. Furthermore, another study, also using HEK-293 cells, transfected with recombinant human TRPV1, calculated the EC50 of CBDV for TRPV1 at 56µM (Starkus *et al*. 2019).

While TRPV1 expression in the central nervous system (CNS) has been controversial (reviewed in Kauer and Gibson 2009), In our model, SVZ-derived cells do not only express high levels of TRPV1, as previously described (Stock *et al*. 2014; Zhai *et al*. 2020; Ramírez-Barrantes *et al*. 2016), but its mRNA expression is significantly increased in the presence of CBDV.

A not well-understood function of TRPV1 is its role in regulating cell death and proliferation. Particularly, the majority of these studies have been carried out using cancer cell models. These highlighted the potential of several TRPV1 modulators, such as Capsaicin, the prototypic TRPV1 agonist, as inducers of cell death (Stock *et al*. 2012; Hou *et al*. 2019; Díaz-Laviada and Rodríguez-Henche 2014), and inhibitors of cell proliferation (Li *et al*. 2021; Lin *et al*. 2013; Weber *et al*. 2016). Thus, ideal candidates as anti-cancer drugs. In our study, we have found that CBDV, in a TRPV1-dependent manner, was able to promote cell viability and proliferation in SVZ-derived NSPCs. Despite that our data contrasts with what was previously shown using cancer cell lines, it aligns with other works that have evaluated the effects of cannabinoids upon cell death and proliferation (Pacher and Mackie 2012; Yang *et al*. 2020; Stone *et al*. 2021; Molina-Holgado *et al*. 2002; Xapelli *et al*. 2013; Rodrigues *et al*. 2017; Bravo-Ferrer *et al*. 2017; Bockmann *et al*. 2022). Interestingly, anandamide, an endogenous cannabinoid and TRPV1 agonist, has been shown to display similar results to the ones obtained in our study (Stock *et al*. 2012; Hofmann *et al*. 2014). In fact, regarding cell death, incubation with anandamide alone did not to induce cell death in the GL261 glioma cell line (Stock *et al*. 2012). In another study, it promoted endothelial cell proliferation in a TRPV1-dependent mechanism of action, since blocking TRPV1 with the antagonist SB366791 inhibited that effect (Hofmann *et al*. 2014). These works support the idea that TRPV1, via cannabinoids, might play a dual role in the mediation of cell death and cell proliferation.

The transmission of electrical signals between neurons in brain networks and circuits is essential to normal brain function. In an oversimplistic model, the neurosphere assay, cultured NSPCs may differentiate into neurons as a consequence of external neural activity that requires a calcium flux (Deisseroth *et al*. 2004). Indeed, our work shows that after a 7-day exposure to CBDV, NSPCs exit the cell cycle and differentiate mostly into neurons. While the mechanisms regarding cell cycle arrest induced by TRPV1 have not yet been fully understood, our results further support the idea that CBDV halts NSPC cell cycle. Thus, the modulation of TRPV1 might offer an innovative strategy to study all these important aspects of well-functioning neural precursors.

Interestingly, our data has shown that CBDV inhibited oligodendroglial maturation and differentiation. Previous works suggested that oligodendrocyte differentiation and myelinating abilities could be modulated by cannabinoids, which are responsible for promoting OPC survival and maturation (Molina-Holgado *et al*. 2002; Rodgers *et al*. 2013). TRPV1 is expressed in OPCs and myelinating oligodendrocytes (Moreno-Luna *et al*. 2021) suggesting that the development of these cells could be modulated by agonists and antagonists of this channel. Therefore, our results agree with published data whilst not reflecting a positive effect on oligodendrogenesis mediated by cannabinoids. The co-incubation with the antagonist 5’-IRTX, was able to reverse the loss of mature oligodendrocytes induced by CBDV. It should, however, be noted that the percentage of myelinated oligodendrocytes in the presence of both drugs is in between that observed in the presence of each drug alone, therefore additive effects cannot be discarded. Importantly, the incubation of 5’-IRTX alone was able to significantly increase the percentage of myelinating oligodendrocytes. One study linked the effect of 5’-IRTX with the inflammatory response induced my TRPV1 on microglial cells both *in vitro* and in samples of cerebrospinal fluid of patients with multiple sclerosis (MS). Upon incubation with 5’-IRTX, the proinflammatory cytokines tumour necrosis factor and interleukin-6, which were elevated after TRPV1 activation by Capsaicin, were significantly reduced (Bassi *et al*. 2019). Our data extends these findings by suggesting that 5’-IRTX not only can have anti-inflammatory properties, but also a pro-oligodendrogenic potential. This response of 5’-IRTX can serve up as a proof of concept for future studies aiming at evaluating the neuroimmune modulatory responses of this drug as well as its pro-oligodendrogenic effects in MS. Further studies, clearly outside the scope of this work, are necessary to better understand the role of CBDV as a modulators of oligodendrocyte differentiation and maturation.

Spontaneous calcium oscillations play important roles in CNS development, neural induction, axon guidance, growth cone morphology, migration, and proliferation (Goswami and Hucho 2007; Komuro and Rakic 1996; Gu *et al*. 1994; Weissman *et al*. 2004). Regarding CBDV, SVZ-derived cells displayed two types of response, one short initial response that corresponds to TRPV1 activation and which is blocked by prior incubation with 5’-IRTX and a long-lasting response that was not fully blocked by the antagonist, possibly suggesting the involvement of a metabotropic receptor. Therefore, the effects in postnatal SVZ neurogenesis might require the activation of both TRPV1 and a putative metabotropic receptor. This polymodal response corresponds to the initial peak of calcium influx observed, prompted by TRPV1, that might act as a trigger for the adjacent cell responses, which are maintained, long term, via a metabotropic mechanism. Thus, when the antagonist 5’-IRTX is present, the trigger is ablated, and the CBDV-elicited calcium response has a delayed onset, as observed.

Furthermore, one study compared the calcium responses elicited by Cannabidiol, to which CBDV is a propyl analogue (Thomas and ElSohly 2016), with the prototypic TRPV1 agonist Capsaicin (Bisogno *et al*. 2001). The calcium influx response evoked by Capsaicin was 100-fold higher than the one elicited by Cannabidiol (Bisogno *et al*. 2001). A recent work highlighted that in cultures derived from rat-dorsal root ganglion neurons, 1µM of Cannabidiol was able to induce a very low calcium response (Anand *et al*. 2020). In our study, CBDV-elicited a calcium response higher than the Cannabidiol reported by Anand and colleagues, and much lower than the one evoked by Capsaicin reported by Bisogno and colleagues. This is relevant since the observed differences regarding the effects of CBDV on cell viability and proliferation might be related to the different magnitudes of calcium influx elicited by different TRPV1 agonists. Indeed, other works have linked the concentration of calcium entering the cell, upon TRPV1 activation, to several molecular pathways that regulate cell homeostasis (Touska *et al*. 2011; reviewed in Zhai *et al*. 2020). The pro-apoptotic effects of TRPV1, as seen in cancer studies, may involve the mitochondria (reviewed in Juárez-Contreras *et al*. 2020). The influx of calcium into the mitochondria results in the depolarization of the mitochondrial membrane and subsequent activation of pro-apoptotic pathways, with higher levels of calcium influx and higher rates of cell death (Kim *et al*. 2006). On the other hand, TRPV1-dependent cell proliferation has also been linked to ATP release and to the activation of the purine receptor P2Y2, with a limited role for calcium influx (Denda *et al*. 2010). Neuronal differentiation has been implicated with calcium influx, with increased levels being linked to a faster differentiation and more mature neurons (Spoerri *et al*. 1990; Holliday *et al*. 1991). Thus, the concentration of calcium influx elicited by CBDV is critical for the maintenance and regulation of cellular homeostasis, with higher concentrations of calcium leading to cell death and inhibition of cell proliferation, as seen for Capsaicin (reviewed in Zhai *et al*. 2020), and lower concentrations promoting cell survival, proliferation and a slower rate of differentiation, as suggested for CBDV (Ramírez-Barrantes *et al*. 2016).

## 5. Conclusions

Based on the multifactorial responses evoked by CBDV, the data herein described provides new insights on the action of this cannabinoid upon TRPV1, with impact in the modulation of postnatal neurogenesis. Despite that the pharmacodynamics of this cannabinoid are still not entirely clear, there is a strong interest by both researchers and clinicians in repurposing CBDV as a viable medicine, with several clinical trials already undergoing studies for several disorders, ranging from autism spectrum disorders (ClinicalTrials.gov Identifier: NCT03537950 2018; ClinicalTrials.gov Identifier: NCT03202303 2017; ClinicalTrials.gov Identifier: NCT03849456 2019) to androgenetic alopecia (ClinicalTrials.gov Identifier: NCT04842383 2021). The fact that CBDV is a non-psychoactive cannabinoid makes this cannabinoid a promising candidate for further studies. Given the significant impact of neurological disorders worldwide, the outcome of such studies has the potential to serve as a cornerstone for advancing brain repair strategies by utilizing NSPCs and cannabinoids as therapeutic options.

### CRediT authorship contribution statement

Conceptualization: DML, AMS, SX; methodology: DML, SSS, SV and SX; investigation: DML, RS, SSS, JM, RSR and JBM; formal analysis: DML and SSS; project administration: AMS, SS and SX; writing – original draft: DML and SX.; writing – review& editing: DML, RS, SSS, JM, RSR, JBM, SV, AMS, SS and SX; funding acquisition: SS, MJD and SX; resources: SV, SS and SX; visualization: DML; supervision: SX.

## Acknowledgements

The authors would like to thank the Bioimaging Unit of Instituto de Medicina Molecular João Lobo Antunes for their technical support. We also acknowledge the funding PPBI-POCI-01-0145-FEDER-022122. We would also like to thank André Gabriel from the ASB Unit of Instituto de Medicina Molecular João Lobo Antunes, for the valuable insight and knowledge on primer design.

## Data availability statement

The data that support the findings of this study are available from the corresponding author upon reasonable request.

## Abbreviations

5’-IRTX: 5’-iodoresinferotoxin

BrdU: 5-bromo-2’-deoxyuridine

CBDV: Cannabidivarin

CB1R: cannabinoid receptor 1

CB2R: cannabinoid receptor 2

DIV: day *in vitro*

ICC: immunocytochemistry

MBP: myelin basic protein

NG2: neuron/glia antigen 2

NSA: neurosphere assay

NSPC: neural stem/progenitor cell

OPC: oligodendrocyte progenitor cell

PI: propidium iodine

qRT-PCR: quantitative real-time reverse transcription polymerase chain reaction

SVZ: subventricular zone

TRPV1: transient receptor potential cation channel subfamily V member 1

NeuN: neuronal nuclei.

## References

Aguirre A., Gallo V. (2004) Postnatal neurogenesis and gliogenesis in the olfactory bulb from NG2-expressing progenitors of the subventricular zone. J. Neurosci. 24, 10530–10541.

Anand U., Jones B., Korchev Y., Bloom S. R., Pacchetti B., Anand P., Sodergren M. H. (2020) CBD Effects on TRPV1 Signaling Pathways in Cultured DRG Neurons. J. Pain Res. 13, 2269–2278.

Anavi-Goffer S., Baillie G., Irving A. J., Gertsch J., Greig I. R., Pertwee R. G., Ross R. A. (2012) Modulation of l-α-Lysophosphatidylinositol/GPR55 Mitogen-activated Protein Kinase (MAPK) Signaling by Cannabinoids. J. Biol. Chem. 287, 91–104.

Arévalo-Martín Á., García-Ovejero D., Rubio-Araiz A., Gómez O., Molina-Holgado F., Molina-Holgado E. (2007) Cannabinoids modulate Olig2 and polysialylated neural cell adhesion molecule expression in the subventricular zone of post-natal rats through cannabinoid receptor 1 and cannabinoid receptor 2. Eur. J. Neurosci. 26, 1548– 1559.

Bassi M. S., Gentile A., Iezzi E., Zagaglia S., Musella A., Simonelli I., Gilio L., et al. (2019) Transient receptor potential vanilloid 1 modulates central inflammation in multiple sclerosis. Front. Neurol. 10, 1–8.

Bisogno T., Hanus L., Petrocellis L. De, Tchilibon S., Ponde D. E., Brandi I., Moriello A. S., Davis J. B., Mechoulam R., Marzo V. Di (2001) Molecular targets for cannabidiol and its synthetic analogues: effect on vanilloid VR1 receptors and on the cellular uptake and enzymatic hydrolysis of anandamide. Br. J. Pharmacol. 134, 845–52.

Bockmann E. C., Brito R., Madeira L. F., da Silva Sampaio L., de Melo Reis R. A., França G. R., da Calaza K. C. (2022) The Role of Cannabinoids in CNS Development: Focus on Proliferation and Cell Death. Springer US.

Bond A. M., Ming G. L., Song H. (2015) Adult Mammalian Neural Stem Cells and Neurogenesis: Five Decades Later. Cell Stem Cell 17, 385–395.

Bravo-Ferrer I., Cuartero M. I., Zarruk J. G., Pradillo J. M., Hurtado O., Romera V. G., Díaz-Alonso J., et al. (2017) Cannabinoid type-2 receptor drives neurogenesis and improves functional outcome after stroke. Stroke 48, 204– 212.

Butti E., Bacigaluppi M., Chaabane L., Ruffini F., Brambilla E., Berera G., Montonati C., Quattrini A., Martino G. (2019) Neural stem cells of the subventricular zone contribute to neuroprotection of the corpus callosum after cuprizone-induced demyelination. J. Neurosci. 39, 5481–5492.

Campbell I. (2007) Chi-squared and Fisher-Irwin tests of two-by-two tables with small sample recommendations. Stat. Med. 26, 3661–3675.

Castaneto M. S., Gorelick D. A., Desrosiers N. A., Hartman R. L., Pirard S., Huestis M. A. (2014) Synthetic cannabinoids: Epidemiology, pharmacodynamics, and clinical implications. Drug Alcohol Depend. 144, 12–41.

Caterina M. J., Schumacher M. A., Tominaga M., Rosen T. A., Levine J. D., Julius D. (1997) The capsaicin receptor: A heat-activated ion channel in the pain pathway. Nature 389, 816–824.

Cavallucci V., Fidaleo M., Pani G. (2016) Neural Stem Cells and Nutrients: Poised Between Quiescence and Exhaustion. Trends Endocrinol. Metab. 27, 756–769.

Cheung T. H., Rando T. A. (2013) Molecular regulation of stem cell quiescence. Nat. Rev. Mol. Cell Biol. 14, 329–340.

ClinicalTrials.gov Identifier: NCT03202303 (2017) Cannabidivarin (CBDV) vs. Placebo in Children With Autism Spectrum Disorder (ASD).

ClinicalTrials.gov Identifier: NCT03537950 (2018) Shifting Brain Excitation-Inhibition Balance in Autism Spectrum Disorder.

ClinicalTrials.gov Identifier: NCT03849456 (2019) Safety and Tolerability of Cannabidivarin (CBDV) in Children and Young Adults With Autism Spectrum Disorder.

ClinicalTrials.gov Identifier: NCT04842383 (2021) Androgenetic Alopecia Treatment Using Varin and Cannabidiol Rich Topical Hemp Oil: a Case Series.

Cohen K., Mama Y., Rosca P., Pinhasov A., Weinstein A. (2020) Chronic Use of Synthetic Cannabinoids Is Associated With Impairment in Working Memory and Mental Flexibility. Front. Psychiatry 11, 1–11.

Compagnucci C., Siena S. Di, Bustamante M. B., Giacomo D. Di, Tommaso M. Di, Maccarrone M., Grimaldi P., Sette C. (2013) Type-1 (CB1) cannabinoid receptor promotes neuronal differentiation and maturation of neural stem cells. PLoS One 8, e54271.

Connor J. P., Stjepanović D., Foll B. Le, Hoch E., Budney A. J., Hall W. D. (2021) Cannabis use and cannabis use disorder. Nat. Rev. Dis. Prim. 7, 1–24.

Costanzo M., Caterino M., Cevenini A., Jung V., Chhuon C., Lipecka J., Fedele R., Guerrera I. C., Ruoppolo M. (2020) Dataset of a comparative proteomics experiment in a methylmalonyl-CoA mutase knockout HEK 293 cell model. Data Br. 33, 106453.

Czaja K., Burns G. A., Ritter R. C. (2008) Capsaicin-induced neuronal death and proliferation of the primary sensory neurons located in the nodose ganglia of adult rats. Neuroscience 154, 621–630.

Deisseroth K., Singla S., Toda H., Monje M., Palmer T. D., Malenka R. C. (2004) Excitation-neurogenesis coupling in adult neural stem/progenitor cells. Neuron 42, 535–552.

Denda S., Denda M., Inoue K., Hibino T. (2010) Glycolic acid induces keratinocyte proliferation in a skin equivalent model via TRPV1 activation. J. Dermatol. Sci. 57, 108–113.

Díaz-Laviada I., Rodríguez-Henche N. (2014) The Potential Antitumor Effects of Capsaicin, in Capsaicin as a Ther. Mol., (Abdel-Salam O. M. E., ed), Vol. 68, pp. 181–208. Springer Basel, Basel.

Ebbert J. O., Scharf E. L., Hurt R. T. (2018) Medical Cannabis. Mayo Clin. Proc. 93, 1842–1847.

Ferreira F. F., Ribeiro F. F., Rodrigues R. S., Sebastião A. M., Xapelli S. (2018) Brain-derived neurotrophic factor (BDNF) role in cannabinoid-mediated neurogenesis. Front. Cell. Neurosci. 12, 1–16.

Figueiredo P. R., Tolomeo S., Steele J. D., Baldacchino A. (2020) Neurocognitive consequences of chronic cannabis use: a systematic review and meta-analysis. Neurosci. Biobehav. Rev. 108, 358–369.

Frey E., Karney-Grobe S., Krolak T., Milbrandt J., DiAntonio A. (2018) TRPV1 agonist, capsaicin, induces axon outgrowth after injury via Ca2+/PKA signaling. eNeuro 5, 1–15.

Galve-Roperh I., Chiurchiù V., Díaz-Alonso J., Bari M., Guzmán M., Maccarrone M. (2013) Cannabinoid receptor signaling in progenitor/stem cell proliferation and differentiation. Prog. Lipid Res. 52, 633–50.

Goswami C., Hucho T. (2007) TRPV1 expression-dependent initiation and regulation of filopodia. J. Neurochem. 103, 1319–1333.

Goswami C., Schmidt H., Hucho F. (2007) TRPV1 at nerve endings regulates growth cone morphology and movement through cytoskeleton reorganization. FEBS J. 274, 760–772.

Grant K. S., Petroff R., Isoherranen N., Stella N., Burbacher T. M. (2018) Cannabis use during pregnancy: Pharmacokinetics and effects on child development. Pharmacol. Ther. 182, 133–151.

Gu X., Olson E. C., Spitzer N. C. (1994) Spontaneous neuronal calcium spikes and waves during early differentiation. J. Neurosci. 14, 6325–6335.

Hall W., Degenhardt L. (2009) Adverse health effects of non-medical cannabis use.

Hill T. D. M. M., Cascio M. G., Romano B., Duncan M., Pertwee R. G., Williams C. M., Whalley B. J., Hill A. J. (2013) Cannabidivarin-rich cannabis extracts are anticonvulsant in mouse and rat via a CB1 receptor-independent mechanism. Br. J. Pharmacol. 170, 679–692.

Hofmann N. A., Barth S., Waldeck-Weiermair M., Klec C., Strunk D., Malli R., Graier W. F. (2014) TRPV1 mediates cellular uptake of anandamide and thus promotes endothelial cell proliferation and network-formation. Biol. Open 3, 1164–1172.

Holliday J., Adams R. J., Sejnowski T. J., Spitzer N. C. (1991) Calcium-induced release of calcium regulates differentiation of cultured spinal neurons. Neuron 7, 787–796.

Hou N., He X., Yang Y., Fu J., Zhang W., Guo Z., Hu Y., et al. (2019) TRPV1 Induced apoptosis of colorectal cancer cells by activating calcineurin-NFAT2-p53 signaling pathway. Biomed Res. Int. 2019, 1–9.

Huizenga M. N., Sepulveda-Rodriguez A., Forcelli P. A. (2019) Preclinical safety and efficacy of cannabidivarin for early life seizures. Neuropharmacology 148, 189–198.

Iannotti F. A., Hill C. L., Leo A., Alhusaini A., Soubrane C., Mazzarella E., Russo E., Whalley B. J., Marzo V. Di, Stephens G. J. (2014) Nonpsychotropic Plant Cannabinoids, Cannabidivarin (CBDV) and Cannabidiol (CBD), Activate and Desensitize Transient Receptor Potential Vanilloid 1 (TRPV1) Channels in Vitro: Potential for the Treatment of Neuronal Hyperexcitability. ACS Chem. Neurosci. 5, 1131–1141.

Juárez-Contreras R., Méndez-Reséndiz K. A., Rosenbaum T., González-Ramírez R., Morales-Lázaro S. L. (2020) TRPV1 channel: A noxious signal transducer that affects mitochondrial function. Int. J. Mol. Sci. 21, 1–17.

Kauer J. A., Gibson H. E. (2009) Hot flash: TRPV channels in the brain. Trends Neurosci. 32, 215–224.

Kee N., Sivalingam S., Boonstra R., Wojtowicz J.. (2002) The utility of Ki-67 and BrdU as proliferative markers of adult neurogenesis. J. Neurosci. Methods 115, 97–105.

Kim S. R., Kim S. U., Oh U., Jin B. K. (2006) Transient Receptor Potential Vanilloid Subtype 1 Mediates Microglial Cell Death In Vivo and In Vitro via Ca 2+-Mediated Mitochondrial Damage and Cytochrome c Release. J. Immunol. 177, 4322–4329.

Komuro H., Rakic P. (1996) Intracellular Ca2+ fluctuations modulate the rate of neuronal migration. Neuron 17, 275– 285.

Kong K. H., Kim H. K., Song K. S., Woo Y. S., Choi W. S., Park H. R., Park M., et al. (2010) Capsaicin impairs proliferation of neural progenitor cells and hippocampal neurogenesis in young mice. J. Toxicol. Environ. Heal. - Part A Curr. Issues 73, 1490–1501.

Laun A. S., Shrader S. H., Brown K. J., Song Z. H. (2019) GPR3, GPR6, and GPR12 as novel molecular targets: their biological functions and interaction with cannabidiol. Acta Pharmacol. Sin. 40, 300–308.

Lecoeur H. (2002) Nuclear Apoptosis Detection by Flow Cytometry: Influence of Endogenous Endonucleases. Exp. Cell Res. 277, 1–14.

Li L., Chen C., Chiang C., Xiao T., Chen Y., Zhao Y., Zheng D. (2021) The Impact of TRPV1 on Cancer Pathogenesis and Therapy: A Systematic Review. Int. J. Biol. Sci. 17, 2034–2049.

Lin C. H., Lu W. C., Wang C. W., Chan Y. C., Chen M. K. (2013) Capsaicin induces cell cycle arrest and apoptosis in human KB cancer cells. BMC Complement. Altern. Med. 13.

Luskin M. B., Boone M. S. (1994) Rate and pattern of migration of lineally-related olfactory bulb interneurons generated postnatally in the subventricular zone of the rat. Chem. Senses 19, 695–714.

MarÉchal E. (2011) Measuring Bioactivity: KI, IC50 and EC50, in Chemogenomics Chem. Genet., pp. 55–65. Springer Berlin Heidelberg, Berlin, Heidelberg.

Marques M. C., Albuquerque I. S., Vaz S. H., Bernardes G. J. L. (2019) Overexpression of Osmosensitive Ca2+-Permeable Channel TMEM63B Promotes Migration in HEK293T Cells. Biochemistry 58, 2861–2866.

Marzo V. Di, Petrocellis L. De (2006) Plant, synthetic, and endogenous cannabinoids in medicine. Annu. Rev. Med. 57, 553–574.

MedCalc Software Ltd (2023) MedCalc’s Comparison of proportions calculator.

Menn B., Garcia-Verdugo J. M., Yaschine C., Gonzalez-Perez O., Rowitch D., Alvarez-Buylla A. (2006) Origin of oligodendrocytes in the subventricular zone of the adult brain. J. Neurosci. 26, 7907–18.

Mishra P., Pandey C., Singh U., Gupta A., Sahu C., Keshri A. (2019) Descriptive statistics and normality tests for statistical data. Ann. Card. Anaesth. 22, 67.

Molina-Holgado E., Vela J. M., Arévalo-Martın A., Almazán G., Molina-Holgado F., Borrell J., Guaza C. (2002) Cannabinoids Promote Oligodendrocyte Progenitor Survival: Involvement of Cannabinoid Receptors and Phosphatidylinositol-3 Kinase/Akt Signaling. J. Neurosci. 22, 9742–9753.

Molina-Holgado F., Rubio-Araiz A., García-Ovejero D., Williams R. J., Moore J. D., Arévalo-Martín Á., Gómez-Torres O., Molina-Holgado E. (2007) CB2 cannabinoid receptors promote mouse neural stem cell proliferation. Eur. J. Neurosci. 25, 629–634.

Moreno-Luna R., Esteban P. F., Paniagua-Torija B., Arevalo-Martin A., Garcia-Ovejero D., Molina-Holgado E. (2021) Heterogeneity of the Endocannabinoid System Between Cerebral Cortex and Spinal Cord Oligodendrocytes. Mol. Neurobiol. 58, 689–702.

Mori H., Ninomiya K., Kino-oka M., Shofuda T., Islam M. O., Yamasaki M., Okano H., Taya M., Kanemura Y. (2006) Effect of neurosphere size on the growth rate of human neural stem/progenitor cells. J. Neurosci. Res. 84, 1682– 1691.

Muller C., Morales P., Reggio P. H. (2019) Cannabinoid ligands targeting TRP channels.

Pacher P., Bátkai S., Kunos G. (2006) The endocannabinoid system as an emerging target of pharmacotherapy. Pharmacol. Rev. 58, 389–462.

Pacher P., Mackie K. (2012) Interplay of cannabinoid 2 (CB2) receptors with nitric oxide synthases, oxidative and nitrative stress, and cell death during remote neurodegeneration. J. Mol. Med. 90, 347–351.

Pertwee R. G. (2005) The therapeutic potential of drugs that target cannabinoid receptors or modulate the tissue levels or actions of endocannabinoids. AAPS J. 7, E625–E654.

Petrocellis L. De, Ligresti A., Moriello A. S., Allarà M., Bisogno T., Petrosino S., Stott C. G., Marzo V. Di (2011) Effects of cannabinoids and cannabinoid-enriched Cannabis extracts on TRP channels and endocannabinoid metabolic enzymes. Br. J. Pharmacol. 163, 1479–1494.

Ramírez-Barrantes R., Cordova C., Poblete H., Muñoz P., Marchant I., Wianny F., Olivero P. (2016) Perspectives of TRPV1 Function on the Neurogenesis and Neural Plasticity. Neural Plast. 2016.

Richardson J. T. E. (2011) The analysis of 2 × 2 contingency tables-Yet again. Stat. Med. 30, 890–890.

Rodgers J. M., Robinson A. P., Miller S. D. (2013) Strategies for protecting oligodendrocytes and enhancing remyelination in multiple sclerosis. Discov. Med. 16, 53–63.

Rodrigues R. S., Ribeiro F. F., Ferreira F., Vaz S. H., Sebastião A. M., Xapelli S. (2017) Interaction between cannabinoid type 1 and type 2 receptors in the modulation of subventricular zone and dentate gyrus neurogenesis. Front. Pharmacol. 8, 1–26.

Rosenthaler S., Pöhn B., Kolmanz C., Huu C. N., Krewenka C., Huber A., Kranner B., Rausch W.-D., Moldzio R. (2014) Differences in receptor binding affinity of several phytocannabinoids do not explain their effects on neural cell cultures. Neurotoxicol. Teratol. 46, 49–56.

Schlag A. K., O’Sullivan S. E., Zafar R. R., Nutt D. J. (2021) Current controversies in medical cannabis: Recent developments in human clinical applications and potential therapeutics. Neuropharmacology 191, 108586.

Soares R., Ribeiro F. F., Lourenço D. M., Rodrigues R. S., Moreira J. B., Sebastião A. M., Morais V. A., Xapelli S. (2020) Isolation and Expansion of Neurospheres from Postnatal (P1-3) Mouse Neurogenic Niches. J. Vis. Exp. 159.

Spoerri P. E., Dozier A. K., Roisen F. J. (1990) Calcium regulation of neuronal differentiation: the role of calcium in GM1-mediated neuritogenesis. Dev. Brain Res. 56, 177–188.

Stanslowsky N., Jahn K., Venneri A., Naujock M., Haase A., Martin U., Frieling H., Wegner F. (2017) Functional effects of cannabinoids during dopaminergic specification of human neural precursors derived from induced pluripotent stem cells. Addict. Biol. 22, 1329–1342.

Starkus J., Jansen C., Shimoda L. M. N., Stokes A. J., Small-Howard A. L., Turner H. (2019) Diverse TRPV1 responses to cannabinoids. Channels (Austin). 13, 172–191.

Stock K., Garthe A., Almeida Sassi F. De, Glass R., Wolf S. A., Kettenmann H. (2014) The capsaicin receptor TRPV1 as a novel modulator of neural precursor cell proliferation. Stem Cells 32, 3183–3195.

Stock K., Kumar J., Synowitz M., Petrosino S., Imperatore R., Smith E. S. J., Wend P., et al. (2012) Neural precursor cells induce cell death of high-grade astrocytomas through stimulation of TRPV1. Nat. Med. 18, 1232–1238.

Stone N. L., England T. J., O’Sullivan S. E. (2021) Protective Effects of Cannabidivarin and Cannabigerol on Cells of the Blood-Brain Barrier under Ischemic Conditions. Cannabis Cannabinoid Res. 6, 315–326.

Straiker A., Wilson S., Corey W., Dvorakova M., Bosquez T., Tracey J., Wilkowski C., Ho K., Wager-miller J., Mackie K. (2021) An evaluation of understudied phytocannabinoids and their effects in two neuronal models. Molecules 26, 1–17.

Suzuki S. O., Goldman J. E. (2003) Multiple cell populations in the early postnatal subventricular zone take distinct migratory pathways: A dynamic study of glial and neuronal progenitor migration. J. Neurosci. 23, 4240–4250.

Thomas B. F., ElSohly M. A. (2016) Biosynthesis and Pharmacology of Phytocannabinoids and Related Chemical Constituents, in Anal. Chem. Cannabis, pp. 27–41. Elsevier.

Thornton C., Dickson K. E., Carty D. R., Ashpole N. M., Willett K. L. (2020) Cannabis constituents reduce seizure behavior in chemically-induced and scn1a-mutant zebrafish. Epilepsy Behav. 110, 107152.

Tominaga M., Tominaga T. (2005) Structure and function of TRPV1. Pflugers Arch. Eur. J. Physiol. 451, 143–150.

Touska F., Marsakova L., Teisinger J., Vlachova V. (2011) A “cute” desensitization of TRPV1. Curr. Pharm. Biotechnol. 12, 122–9.

Urbán N., Blomfield I. M., Guillemot F. (2019) Quiescence of Adult Mammalian Neural Stem Cells: A Highly Regulated Rest. Neuron 104, 834–848.

Vilain S., Esposito G., Haddad D., Schaap O., Dobreva M. P., Vos M., van Meensel S., Morais V. A., de Strooper B., Verstreken P. (2012) The yeast complex I equivalent NADH dehydrogenase rescues pink1 mutants. PLoS Genet. 8.

Wahl P., Foged C., Tullin S., Thomsen C. (2001) Iodo-Resiniferatoxin, a New Potent Vanilloid Receptor Antagonist. Mol. Pharmacol. 59, 9–15.

Weber L. V., Al-Refae K., Wölk G., Bonatz G., Altmüller J., Becker C., Gisselmann G., Hatt H. (2016) Expression and functionality of TRPV1 in breast cancer cells. Breast Cancer Targets Ther. 8, 243–252.

Weissman T. A., Riquelme P. A., Ivic L., Flint A. C., Kriegstein A. R. (2004) Calcium waves propagate through radial glial cells and modulate proliferation in the developing neocortex. Neuron 43, 647–661.

Xapelli S., Agasse F., Sardà-Arroyo L., Bernardino L., Santos T., Ribeiro F. F., Valero J., et al. (2013) Activation of Type 1 Cannabinoid Receptor (CB1R) Promotes Neurogenesis in Murine Subventricular Zone Cell Cultures. PLoS One 8, e63529.

Yang S., Hu B., Wang Z., Zhang C., Jiao H., Mao Z., Wei L., Jia J., Zhao J. (2020) Cannabinoid CB1 receptor agonist ACEA alleviates brain ischemia/reperfusion injury via CB1–Drp1 pathway. Cell Death Discov. 6.

Zhai K., Liskova A., Kubatka P., Büsselberg D. (2020) Calcium entry through trpv1: A potential target for the regulation of proliferation and apoptosis in cancerous and healthy cells. Int. J. Mol. Sci. 21, 1–25.

Zimmermann T., Maroso M., Beer A., Baddenhausen S., Ludewig S., Fan W., Vennin C., et al. (2018) Neural stem cell lineage-specific cannabinoid type-1 receptor regulates neurogenesis and plasticity in the adult mouse hippocampus. Cereb. cortex 28, 4454–4471.

