## Supplementary material for "Unravelling a novel role for Cannabidivarin in the modulation of subventricular zone postnatal neurogenesis": Fig. S1; Table S1

### Supplementary Figures

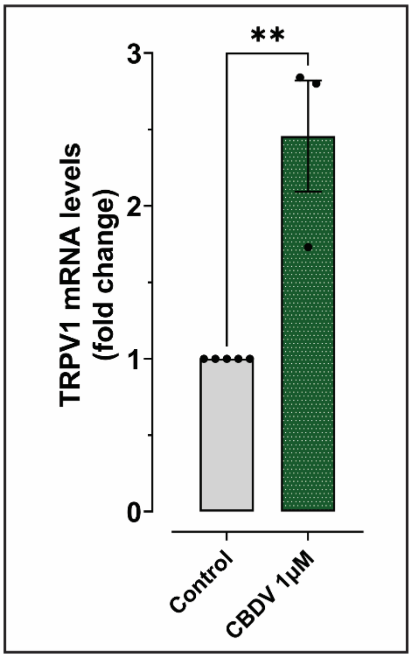


**Fig. S1 – TRPV1 expression is increased by Cannabidivarin under differentiative conditions.** CBDV treatment at DIV7 promoted an increase in TRPV1 mRNA expression in SVZ-derived cells. **(A)** Bar graph depicts the levels of TRPV1 mRNA expression. Values were normalized to the control expression of GAPDH expression for each experiment. Data presented as Mean ± SEM and the control was set to 1. n=3-5; **p<0.01. Unpaired t test.

### Supplementary Tables

**Table S1 – Detailed numbers and percentages of neurospheres analysed distributed according to diameter ($D_N$) (related to Figure 3).**

| | | Control | CBDV 1μM |
|---|---|---|---|
| **30μm < $D_N$ ≤ 100μm** | **n** | 1177 | 856 |
| | **%** | 54.07 | 47.77 |
| **100μm < $D_N$ ≤ 300μm** | **n** | 970 | 872 |
| | **%** | 44.56 | 48.66 |
| **300μm < $D_N$ ≤ 500μm** | **n** | 21 | 51 |
| | **%** | 0.96 | 2.85 |
| **500μm < $D_N$ ≤ 700μm** | **n** | 7 | 8 |
| | **%** | 0.32 | 0.45 |
| **$D_N$ > 700μm** | **n** | 2 | 5 |
| | **%** | 0.09 | 0.28 |
| **Total** | **n** | 2177 | 1792 |
| | **%** | 100 | 100 |
